# Clonal evolutionary analysis reveals patterns of malignant transformation in pancreatic cancer from Intraductal Papillary Mucinous IPMN Neoplasms (IPMN)

**DOI:** 10.1101/2024.08.02.606217

**Authors:** Antonio Pea, Xiaotong He, Rosie Upstill-Goddard, Claudio Luchini, Leonor Patricia Schubert Santana, Stephan Dreyer, Fraser Duthie, Nigel B. Jamieson, Colin J. McKay, Euan J. Dickson, Alessandra Pulvirenti, Selma Rebus, Genomics Innovation Alliance, Scottish Genome Partnership, Paola Piccoli, Nicola Sperandio, Rita T. Lawlor, Michele Milella, Fieke E. M. Froeling, Roberto Salvia, Aldo Scarpa, Andrew V. Bainkin, David C. Wedge, David K. Chang

## Abstract

Intraductal papillary mucinous neoplasms (IPMNs) are critical precursors to pancreatic ductal adenocarcinoma (PDAC), a highly lethal cancer due to late detection and rapid progression. Using multi-region whole-genome and transcriptome sequencing, we traced the evolution of PDAC from IPMN, constructing detailed phylogenetic trees to provide insights into subclonal architectures and progression pathways. Our analysis identified two distinct evolutionary trajectories: one driven by a single ancestral clone, and another involving multiple independent ancestral clones, potentially influencing the timing and nature of PDAC onset. We further explored the roles of mutational signatures and structural variants (SVs) in promoting clonal evolution. Complementing these genomic findings, our transcriptomic analysis revealed unique gene expression profiles and variations in the immune landscape, correlating with the different progression stages from IPMN to PDAC. These insights reveal the complex molecular dynamics of IPMN progression to PDAC, highlighting the need to refine early detection and treatment strategies.

## INTRODUCTION

Pancreatic ductal adenocarcinoma (PDAC) is associated with a dismal prognosis and is projected to be the second leading cause of cancer related deaths soon(1). This poor prognosis is largely attributed to the late-stage presentation, which is further compounded by rapid disease progression once diagnosed.

As one of the precursor lesions of pancreatic cancer, intraductal papillary mucinous neoplasm (IPMN) are macroscopic cystic lesions that are radiologically detectable offering a window of opportunity for detection, and intervention to prevent malignant transformation. Current clinical recommendations for surgical resection of IPMNs are primarily based on imaging criteria such as size, leading to large numbers of patients undergoing annual imaging surveillance(2). A better understanding of the critical molecular events that drive progression in premalignant lesions is urgently needed to better manage these patients to achieve early diagnosis and cancer prevention.

Recent genomic studies on PDAC have mainly focused on a single region of each cancer(3–6). without detailed pathology and radiology annotations leading to cohort genomics studies of PDAC that have arisen from a mixture of all types of precursor lesions. Malignant transformation from dysplasia to invasive cancer in the early stages of cancer evolution has been modelled using single driver mutations to define the relationship between precursor lesions and invasive cancer(7–9),(10). However, IPMN intratumoral heterogeneity and tumour evolution have not been characterised using multi-regional samples from dysplasia and cancers from the same patients at whole genome resolution to date.

Here, we performed multi-region whole genome and transcriptome sequencing of non-invasive IPMNs and their histologically associated PDACs. We clustered genomic alterations, constructed phylogenetic trees, and determined the mutational signatures for multiple samples within each tumour to gain insight into the subclonal architecture and the clonal dynamics during the progression from IPMN to PDAC. We carried out integrative analysis of WGS and RNAseq data based on gene expression and cell type deconvolution, demonstrating distinct tumour evolutionary trajectories and distinctive footprints of genomic evolution, and identified key functional signalling networks involved in the progression from non-invasive IPMN to PDAC.

## RESULTS

### Patient characteristics

A cohort of 12 patients underwent pancreatic resection for IPMNs deemed at high clinical risk according to clinical guidelines. The detailed clinicopathological features of the patient cohort are presented in Table S1. Forty-seven tumour samples were harvested from these 12 surgical specimens at a mean of 4 tumour regions (range 2-6) each, plus a matching normal sample from each patient. From these, we performed whole-genome sequencing (WGS) on 54 samples (42 tumour samples, mean coverage 80X, and 12 normal samples at 60X). Of these, 37 tumour samples were included in the clonal analysis, after excluding 5 samples based on low estimates of tumour purity or total mutation number. Transcriptome sequencing (RNAseq; average 100 million paired reads) was performed on 36 samples, of which 32 had matching WGS data from the same sample. A study sample CONSORT diagram is presented in supplementary material.

### Mutational landscape of IPMN to PDAC malignant transformation

We identified 66,724 SNVs, 4,683 Indels and 2,447 structural variants (SVs) across all tumour samples. SNV, Indel and SV events were seen in 48 driver genes and 11,200 other genes across IPMN and associated PDAC samples. We observed significantly higher mean tumour mutation burden (TMB) (4.11 vs 1.38, *p* = 0.0001; Figure S1B1a, Table S1) and mean number of SNVs (3,558 vs 1,608, *p* = 0.0022; TableS1), Indels (253 vs 105.5, *p* = 0.0088; TableS1) and SVs (91.8 vs 31.7, *p* = 0.0006; TableS2) in PDAC than in IPMN samples. These differences were consistently seen in coding and non-coding mutations, and in both drivers and passengers (Figure 1B.a-c). When SVs were sub-classified as large fragment translocations, tandem duplications, deletions or inversions, the average number of each SV type was also significantly higher in PDAC than in IPMN samples (Figure S1B1c and Table S2). Driver genes of interest were identified as missense and nonsense SNVs, and as frameshift deletions / insertions and essential splice site deletions. Across these categories of SNVs and Indels, mutations were detected in 17 driver genes, comprised of eight tumour suppressors (*ARID1A, ATM, CDKN2A, LRP1B, NBEA, RNF43, SMAD4, TP53)* and five oncogenes (*FAT3, GNAS, KMT2C, KRAS, ZNF521).* In line with previous studies, *KRAS* was the most frequently mutated gene (22/41), occurring in both IPMN and PDAC samples (Figure 1C). The most common mutation in *KRAS* was G12D (15/22), followed by G12V (5/22) then G12R (2/22). Two cases (Case 4 and 15) harboured two different variants of *KRAS*, namely G12D-G12V and G12V-G12R (Figure S1C2, Table S3), However, significant differences of the *KRAS* cancer cell fractions (CCFs) were not found between IPMN and PDAC samples in our study cohort (Figure S1C3). As expected SNVs and Indels of *TP53* were seen in IPMN with HGD and in PDAC samples. *LRP1B* mutation was significantly associated with invasive cancer (0 in IPMN vs 7 in PDAC, *p* = 0.0056), whereas GNAS mutation was significantly associated with non-invasive low grade dysplasia (8 in IPMN vs 1 in PDAC, *p* = 0.0297) (Figure S1C1). Driver SVs in exonic regions were detected in Case 7 and Case 16, including cases with *MUC4* in high grade IPMN and PDAC, *MDM2* in both low- and high-grade lesions, and *MYB, THRAP3, PDCD1LG2,* in high grade IPMN and PDAC lesions (Figure 1C).

**Fig. 1.**
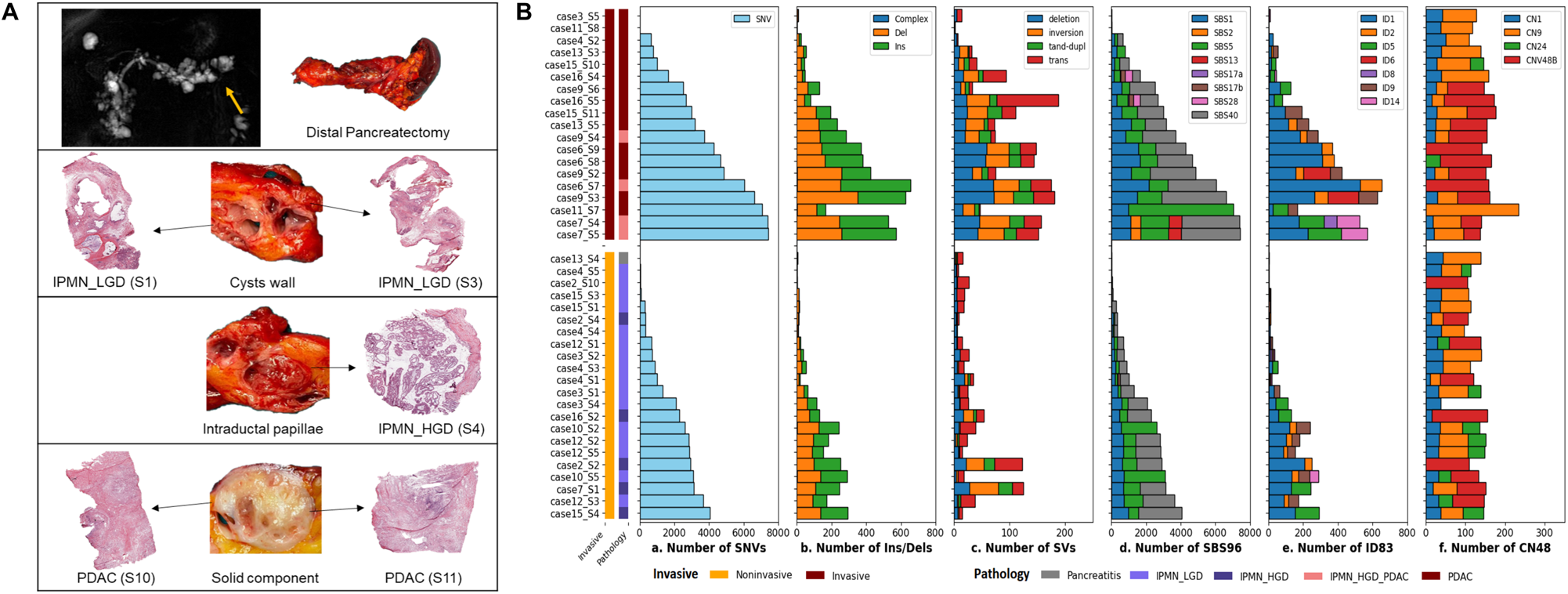
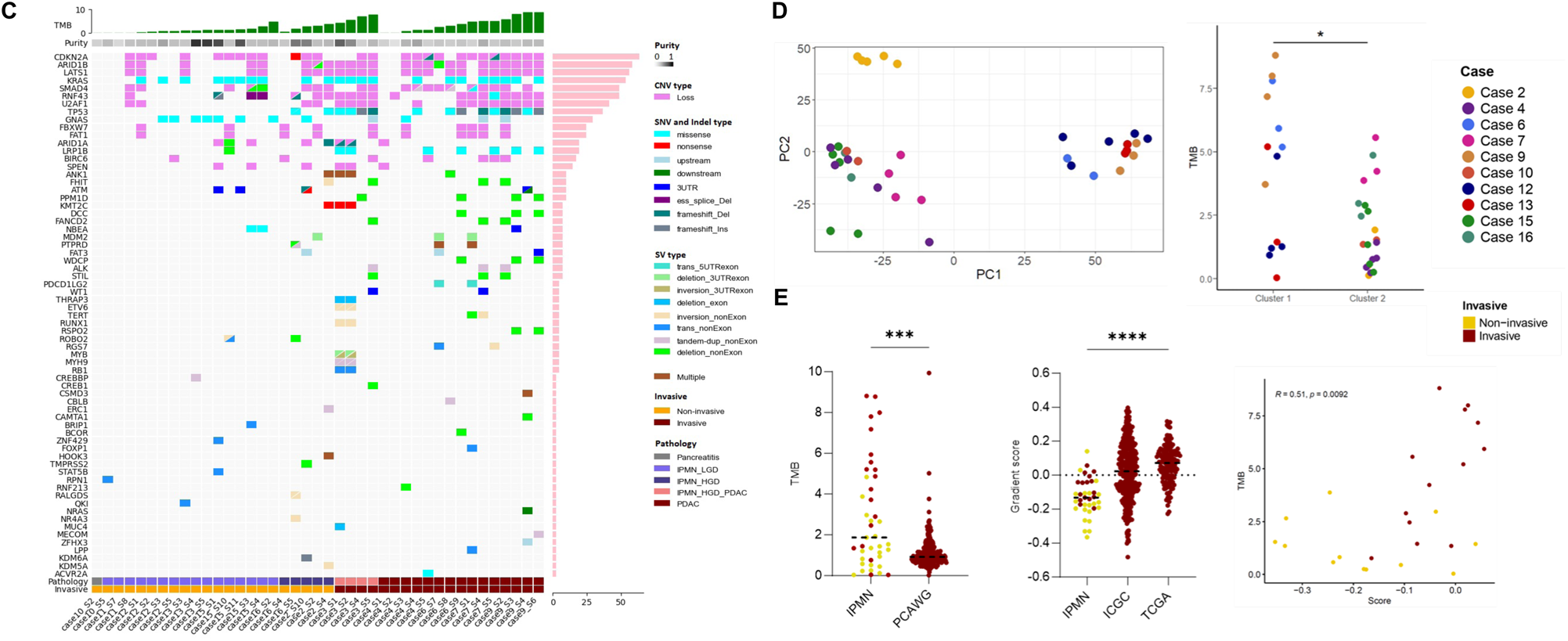
**A) Multiregional Sampling Protocol.** Case 15 is presented as a representative example of our multiregional sampling approach, designed to capture samples across different grades of dysplasia and invasive cancer within a single neoplastic lesion. Preoperative MRI for Case 15 revealed a multifocal IPMN in the pancreas with a solid component in the tail (yellow arrow), necessitating a distal pancreatectomy. Fresh tissue samples were procured for downstream molecular analysis from the walls of various cysts revealing IPMN with low grade of dysplasia (IPMN_LGD), intraductal papillae revealing an IPMN with high grade of dysplasia (IPMN_HGD), and the adjacent solid component revealing pancreatic ductal adenocarcinoma (PDAC). **B) Mutational landscape of IPMNs and PDACs**. Total number of SNVs (a), INDELS (b), SVs (c), including deletion, inversion, tandem duplication and translocation, in each sample, grouped as IPMN (Lower) and PDAC (Upper). The average number of SNVs, INDELS and the average number of each breakpoint type was significantly increased in PDAC compared to IPMN (p < 0.05 for each genomic alteration). pIPMN-HGD and PDAC compared to that in IPMN-LGD. E, Total number and proportions of Indel (ID83) COSMIC signatures in each sample. 7 ID signatures including ID1, 2, 5, 6, 8, 9, and 14 were identified in 41 samples. F, Total number and proportion of copy number (CN48) COSMIC signatures in each sample. 4 CN signatures including CN1,9,24, and a novel CN signature CNV48B were identified in 41 samples. **C) Driver SNVs, INDELs, SVs and CNVs of IPMNs and PDACs**. Coding driver SNVs and INDELs in 41 samples. Drivers of interest were identified in SNVs with missense, nonsense and silent while Indels with del-frameshift, Ins-frameshift and dell_ess splice. 13 mutational drivers were detected in all samples comprising 7 tumour suppressors ARID1A, CDKN2A, LRP1B, NBEA, RNF43, SMAD4, TP53 and 6 oncogenes ATM, FAT3, GNAS, KMT2C, KRAS, ZNF521. KRAS missense was the most frequently mutated (22/34) and constantly occurred from IPMN to PDAC. TP53 and LRP1B mutation showed a significant association with high-grade lesions (p < 0.01) whereas GNAS mutations were associated with the low-grade lesions (p<0.05). 47 drivers were involved in SVs including translocation, tandem duplication, deletion, and inversion. Driver breakpoints in exon region were detected in case 7 and case16, including MUC4 in low grade lesion IPMN-PDAC, MDM2 in both low-high grade lesions, and MYB, PDCD1LG2, THRAP3 in PDAC. Chromosomal rearrangements in of drivers were found predominantly in the non-exon regions. CNVs in 13 different cytobands were significantly associated with IPMN-PDAC progression including 7 gains and 5 losses. Copy number losses were identified in 11 drivers and the most common alteration is CDKN1A, followed by LAT1S1, ARID1B, SMAD4, U2AF1, RNF43, FBXW7, BIRC6, SPEN, ARID1A. Losses of RNF43 and U2AF1 were significantly associated with high-grade lesions (p < 0.05). **D) PCA of gene expression, revealing two distinct clusters.** Cluster 1 (cases 6, 9, 12, 13) and Cluster 2 (cases 2, 4, 7, 10, 15, 16). Comparison of TMB based on SNV-indel between the two identified groups. Cluster 1 is characterized by higher TMB (p=0.02). **E) TMB and the transcriptomic gradient score comparison between IPMN and PDAC. a.** TMB in the IPMN cohort (41 samples) and in the PCAWG cohort (241 samples). **b.** Distribution of transcriptomic gradient scores (classical vs. squamous) in the 37 study samples, compared to 362 ICGC and 149 TCGA cases, with higher scores indicating more squamous tumor. **c.** Scatter plot showing the correlation between TMB and the gradient score within the IPMN cohort. Lower transcriptomic scores in IPMNs suggest an earlier disease stage, with a trend towards higher TMB and increasing squamous characteristics as the disease progresses.

Copy number alterations (CNAs) have been shown to contribute to cancer initiation and progression(11), informing mechanisms of tumour progression. Using Battenberg and GISTIC2(12), we examined the chromosomal regions of CNA with significantly high frequency in 41 tumour-normal pairs. High frequency CNAs were identified in 64 regions (33 gains, 18 losses and 13 complex regions). The most frequently aberrant CNA region was 17p11.2 complex (gain 13/41, loss 12/41, complex 15/41) followed by 20q11.21 gain (36/41) and 9p21.3 loss (25/41). Most of the CNAs (52/64) were carried through from non-invasive IPMN to PDAC (Figure S2A), with four gains and four losses. having higher prevalence in PDAC than in IPMN samples. Intriguingly, one gain at 17p11 had significantly higher prevalence in IPMN than PDAC samples(Figure S2B, Table S4). Furthermore, CN losses were identified in 11 drivers, the most frequent being *CDKN2A* (9q21.3, 25/41), followed by *LAT1S1* (6q27, 23/41), *ARID1B* (6q27, 23/41), *SMAD4* (18q21.2, 19/41), *U2AF1* (21q22.3), *RNF43* (17q22), *FBXW7* (4q34.3), *BIRC6* (2p22.1), *SPEN* (1p36.13), *ARID1A* (1p36.13), with large region losses covering *RNF43* and *U2AF1* being significantly associated with high-grade IPMN (Figure S1C1, *p* = 0.0086 and *p* = 0.0049).

### Mutational signatures in IPMN and PDAC

Mutational signature analysis extracted 8 single-base substitution (SBS) COSMIC signatures (Figure 1B), of which SBS1 and SBS5 (clock-like / age) were the most frequent (39/41 samples) and were observed in samples from all cases. The rest of the SBS signatures are largely of unknown aetiology with the second most common being SBS40 (35/41 samples; in 9 out of 10 cases). The rare SBS signatures SBS17a and SBS17b were present in Cases 4 and 16 and SBS28 was only seen in Case 16. Of interest, the AID / APOBEC family associated signatures SBS2 and SBS13(13) were only seen in Case 7, and only in those clones that progressed to invasive cancer (Figure 1B.d and FigureS1B2a). Indel signatures ID1 and ID2, shown to be associated with DNA mismatch repair deficiency(14), were seen in both low- and high-grade lesions of 12 and 7 cases respectively. In addition, 5 other Indel signatures with unknown aetiology were detected across IPMN and PDAC samples including ID9 (21/41), ID5 (13/41), ID14 (10/41), ID8 (4/41) and ID6 (2/41) (Figure 1B.e and Figure S1B2b). Copy number signature CN9, caused by chromosomal instability and associated with poor survival in various cancer types(15), was seen in 31 out of 41 IPMN and PDAC samples from 11 cases. In addition, a novel CN signature CN48B with unknown mechanism was extracted in 21 samples from 10 cases (Figure 1B.f and Figure S1B2c). This previously unreported signature is characterised by a large number of LOH segments of varying lengths, which are predominantly distributed across high copy numbers (Figure S1B3).

### Heterogeneity in genomic evolution of IPMN to PDAC

To further understand the origin of the malignant subclone and to map comprehensively the genomic evolution from pre-malignant IPMN to invasive PDAC, we clustered mutations according to their CCFs(16) and constructed phylogenetic trees from multi-regional samples from 11 cases (Figure 2A). From the hierarchy of oncogenic events, we constructed phylogenies representing the genetic relationship between all subclones found in IPMN and PDAC samples. These phylogenies were characterised by two distinctive trajectories of genomic evolution in IPMN to PDAC progression: (1) all IPMNs and PDACs develop from one single most recent common ancestor (MRCA) (N = 8; Cases 3, 6, 7, 9, 10, 12, 13, 16), and (2) IPMNs and PDACs develop from separate, independent MRCAs (N = 3; Cases 2, 4 and 15).

**Fig. 2.**
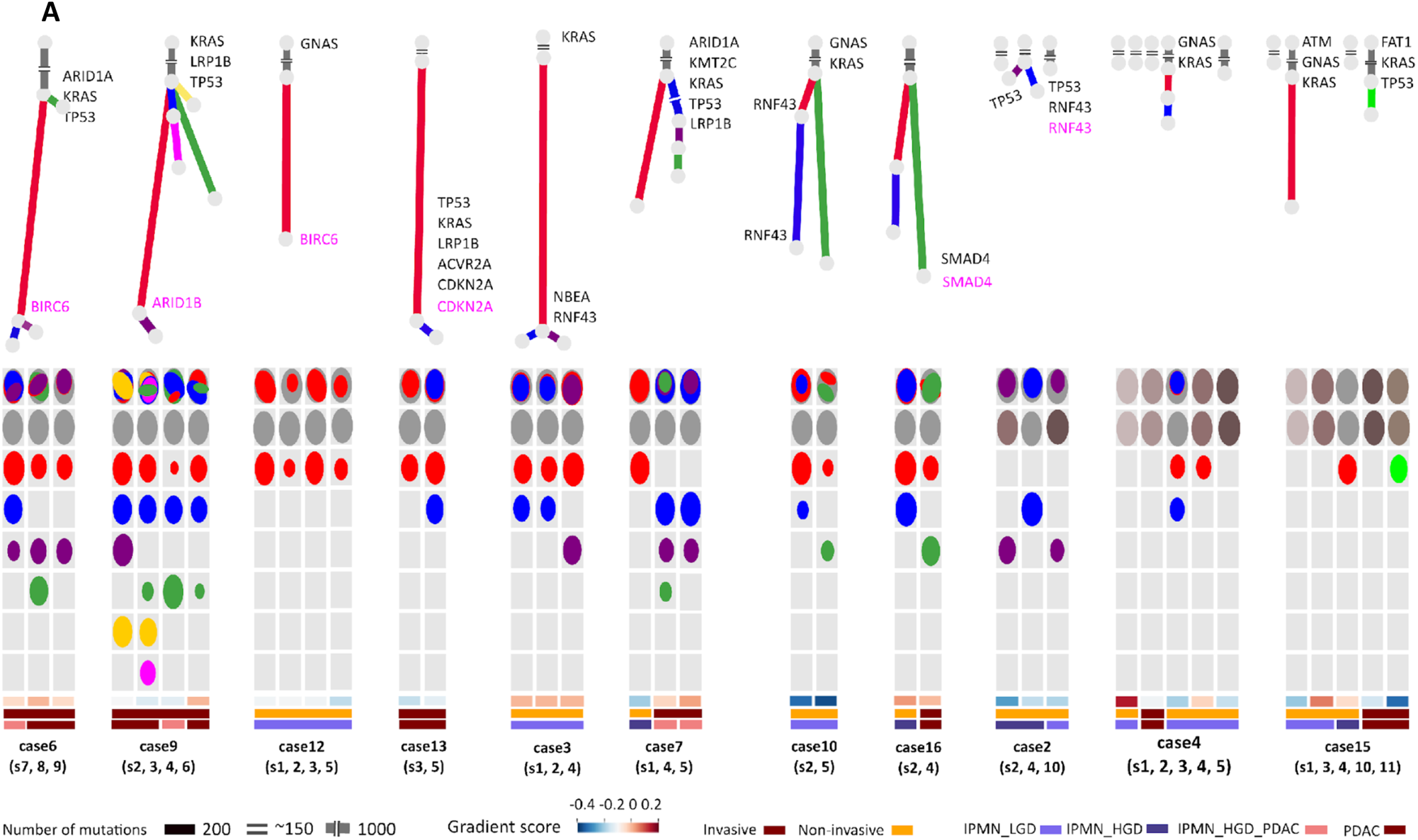
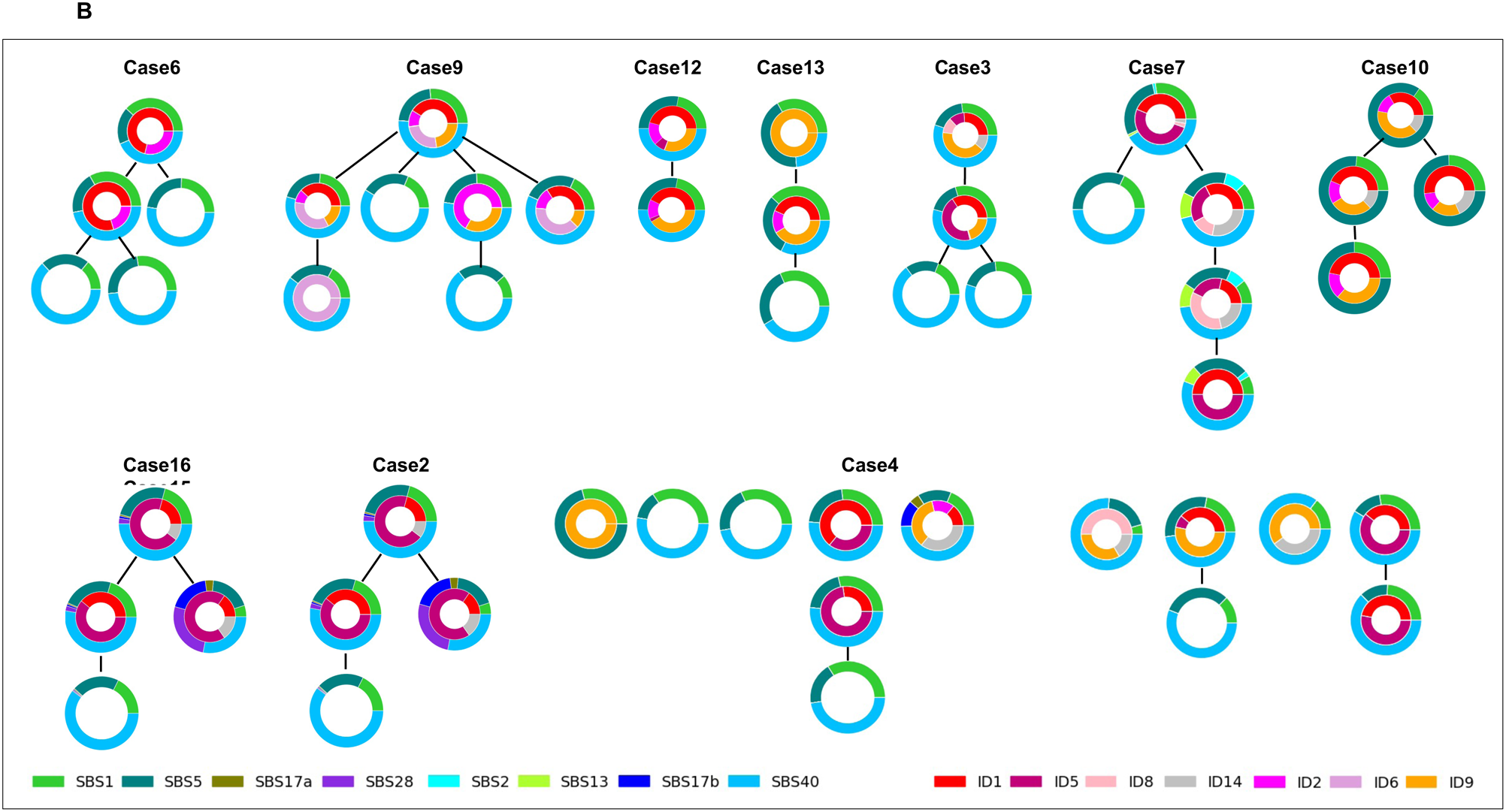
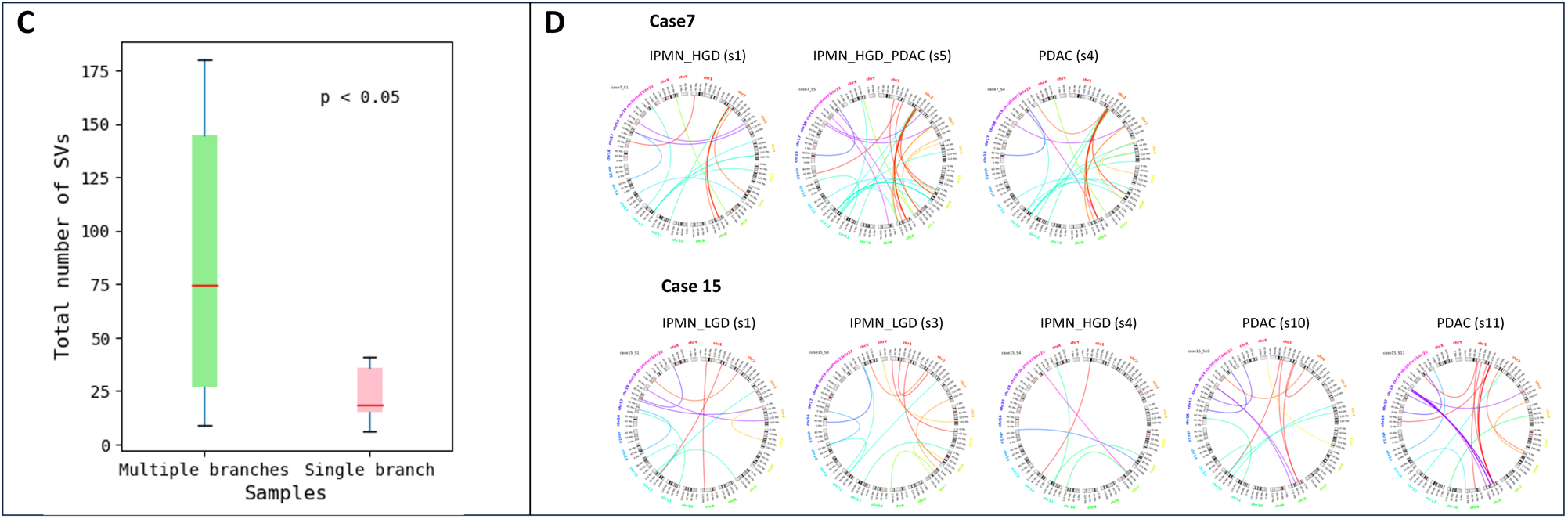
Patterns of genomic heterogeneity and evolution in 11 cases of IPMN-PDACs. **A, Subclone structure within IPMNs and PDACs.** All the subclones identified in the WGS samples are reconstructed as phylogenetic trees (Upper) and oval plots (Lower). Phylogenetic trees show the relationships between subclones in the samples of each patient. Branch lengths are proportional to the number of SNVs and INDELs in each cluster. Branches are annotated with samples in which they are presented and with SNV/INDEL (Black) or CNV (Pink) assigned to that subclone. On the left side, the subclonal seeding in the 8 cases fully depend on a common MRCA. On the right side, the tumour clones in the 3 cases are initiated from multiple independent tumours. The ovals in different colour indicate each subclone identified across multiple samples from each patient (Supplementary Table S7). Each row represents a sample, with ovals in the first row nested if required by the pigeonhole principle as described in Methods. The area of the ovals is proportional to the CCF of the corresponding subclone. **B, Mutational Processes in Clones and Subclones of IPMN-PDACs**. The phylogenetic trees are diagrammed with unscaled branches in which they are annotated with mutational signatures. As shown in each pie chart, the clonal-specific signatures are integrated into individual branches in cancer cell phylogenies. The color-coded area of the pie chart is proportional to the SBS96 or ID83 signature of the corresponding SNVs or INDELs in each branch. The most common signatures SBS1, 5 and 40 are identified in both single MRCA and monoclonal origins and exist in either clone or subclone. In the most of cases (2, 3, 4, 6, 7, 9, 15, 16), ID1 and/or ID2 join the early stages of tumour initiation but are not present throughout all subclonal seedings. **C, SV comparison between case-sample with multiple branches and single branches.** Cases with branching clonal evolution showed a higher count of structural variations. **D, Structural Variations in Clonal Evolution of IPMN Cases.** Case 7, characterized by single-branch clonal evolution, shows PDAC samples with increased SVs from HGD, alongside new SVs in HGD, indicating ongoing evolution after the appearance of the infiltrating clone. Case 15, demonstrating multi-branch clonal evolution, features distinct SVs patterns in samples, each derived from independent ancestral clones, underscoring their independent development.

Further heterogeneity in tumour evolution was also observed within both single and multiple independent MRCA trajectories. For the single MRCA cases, driver alterations occurred both clonally and subclonally, including driver SNVs, Indels and CNAs in both oncogenes and tumour suppressor genes in Cases 3, 6, 7, 9, 10 and 12. However, in Cases 13 and 16, driver alterations were only seen in subclones. For the multiple independent MRCA cases, within the multiple samples of Case 15, two separate tumours were initiated from different MRCAs that both harboured driver mutations. However in other clones without driver mutation, they behaved inactively in subclonal seeding. This same phenomenon was also seen in the independent tumour samples of Cases 2 and 4. Uneven branch lengths in the trees were observed in Cases 6, 7, 9 and 10. The most striking example is Case 6, characterised by 1,932 mutations in the red branch and only 32 in the green branch, suggesting a subclonal sweep occurring on the red branch much later than that on the green branch (Figure 2A). These results suggest critical genomic events not only happened in the MRCAs of our tumours but also happened in descendant clones, contributing to malignant transformation.

### Single vs multiple branch clonal evolution in IPMN

By investigating whether lineages in different patients are associated with specific genomic alterations, we observed that expansion of subclones within IPMN can follow one of two different patterns: (1) the ancestral clone gives rise to a single subclone, or (2) the ancestral clone branches into multiple subclones that diverge independently from each other.

In the linear model of clonal progression, a dominant clone acquires successive alterations, generating a linear series of subclones. Each newly formed subclone in this model retains the genetic traits of its predecessor while gaining its own unique alterations. This pattern suggests a stepwise progression of disease, with each evolutionary ‘step’ marked by the emergence of a new subclone. Cases 4, 12, 13 and 15 exemplify this linear clonal evolution. The second pattern was observed in Cases 2, 6, 7, 9, 10, and 16 and is characterised by multiple subclones diverging independently from the original or ancestral clone, creating a complex clonal architecture with multiple branches. Each of these branches represents a distinct subclonal lineage, harbouring unique genetic alterations in addition to those of the ancestral clone. A caveat to the separation of tumours by their branching or linear pattern is that the observed tree shape will be affected by random sampling of regions of IPMN and PDAC. However, we noted that cases displaying branching clonal evolution harbour significantly higher levels of genomic instability compared to those displaying the linear pattern. A significantly higher count of intrachromosomal structural variations (mean of 85.2 vs 29.7, *p = 0.009*) was indeed observed in these cases, suggesting that SVs might be contributing to the generation of diverging subclones (Figure 2D).

### Temporal dynamics of mutational signatures

Cancer genomes evolve continuously due to the accumulation of mutations, and in turn mutational processes change over time, leaving dynamic signatures in the accumulated genomic variation in each tumour(17). To better understand the molecular events that drive malignant transformation, we detailed the changes of the mutational processes during tumour evolution from IPMN to PDAC, by calculating the proportion of mutations attributed to each mutational signature separately for each of the branches within the tumour phylogenies (Figure 2B). Overall, SBS1, 5 and 40 were the most prevalent signatures and were identified in both single MRCA and multiple MRCA trajectory cases as well as in both trunks and branches of the phylogenies. The AID / APOBEC family associated signatures SBS2 and SBS13 emerged in the MRCA of Case 7 but remained active only on the branch associated with invasive PDAC. Similarly, in Case 16, there was a distinctive burden of SBS28 in the invasive PDAC specific subclone and not in the pre-malignant IPMN branch. ID1 and / or ID2 signatures, known to be related to DNA mismatch repair deficiency, were seen in the early stages of tumour initiation in most of the cases (2, 3, 4, 6, 7, 9, 15, 16); however they were unexpectedly absent in all subclones. A similar phenomenon was also observed in some cases harbouring ID5 (unknown), ID6 (HR deficiency) and ID9 (unknown) signatures in earlier clones but not in subclones. A caveat to our analysis is that the number of mutations assigned to many of the subclones was much lower than that assigned to the trunk, and there were much fewer indels than SNVs in all samples, raising the possibility of non-detection of an indel signature due to insufficient power.

### Modelling malignant transformation from IPMN to PDAC

Out of the 12 cases subjected to multi-regional sampling and clonal evolutionary analysis, six distinct cases provided samples representing different grades of dysplasia and histologically associated cancer within the same surgical specimen. This includes two cases (Case 6 and 9) with samples containing IPMN with microinvasive PDAC (IPMN_HGD_PDAC). Four other cases (Case 7, 4, 15, 16) provided samples of both IPMN and invasive cancer. In one other case (Case 2), even though we collected and performed WGS on both non-invasive IPMN and PDAC samples, WGS analysis was successful only for IPMN samples and not the invasive PDAC samples.

For each of these six cases, the branch point on the phylogenetic tree where IPMN and PDAC diverge is indicative of the time of emergence of the infiltrative clone(s). The clonal architecture in Case 9 reveals a branching evolutionary pattern, marked by an ancestral clone (grey) and several independent subclones (Figure 2A). Invasive cancer initiation is attributed to the ubiquitous clone, present across all samples, possessing driver mutations in *KRAS*, *TP53*, and *LRP1B*. Its presence in both HGD IPMN and microinvasive PDAC indicates infiltrative potential at an early stage. The independent “blue” and “green” subclones, detected in both HGD IPMN and PDAC samples, seem also to display invasive potential, whereas the emergence of other subclones confined to the PDAC samples (yellow, pink and purple subclones) suggest a secondary wave of clonal expansion. One PDAC sample (S3) exhibited multiple intrachromosomal SVs and included most of the subclones, suggesting that the intrachromosomal SVs may have driven subclone development in this case.

Similarly, in Cases 6 and 16, a single MRCA is ancestral to all the HGD and the PDAC samples, suggesting that the MRCA present in HGD may have already acquired invasive capabilities. In Case 6, a similar pattern of SVs was observed in both HGD and PDAC samples. Most notably, PDAC samples (S4 and S5) in Case 16 demonstrated a higher number of intrachromosomal SVs compared to the HGD sample (S2), which may be the drivers for invasive disease and the drivers of PDAC specific subclones (green). In Cases 4 and 15, independent ancestral clones were identified in IPMN and invasive cancer samples. Specifically, in Case 15, unique clones with mutations in *KRAS*, *GNAS*, and *ATM were* observed in HGD, while mutations in *KRAS* and *TP53* were seen in invasive cancer clones (Figure 2A). This suggests the presence of independent evolutionary trajectories for the HGD and invasive cancer clones within the same patient. These observations demonstrate the possibility of multiple oncogenic events occurring within distinct cell populations that lead to independent clonal expansions and parallel evolution, possibly occurring at different timepoints of the disease progression.

### Integration of multi-omics profiles with phylogenetic patterns in IPMN

Principal Component Analysis (PCA) of RNAseq from samples matched to those used for WGS identified two gene expression clusters: Cluster 1 (Cases 6, 9, 12, 13) and Cluster 2 (Cases 2, 4, 7, 10, 15, 16) (Figure 1D). Cluster 1 exhibited lower expression of most of the cancer hallmarks, except those related to development, bile acid metabolism, and downregulated *KRAS* signalling(18), and samples in this cluster exhibited higher expression of the genes *TP53*, *S100A2*, *KRT6A* and *LRP1B* (Figure 3A, 3B). Genomically, Cluster 1 was characterised by higher TMB (combining SNV and indels; *p* = 0.02) and enriched for mutations in *LRP1B* (Figure 3C), with no discernible differences in histology.

**Fig. 3.**
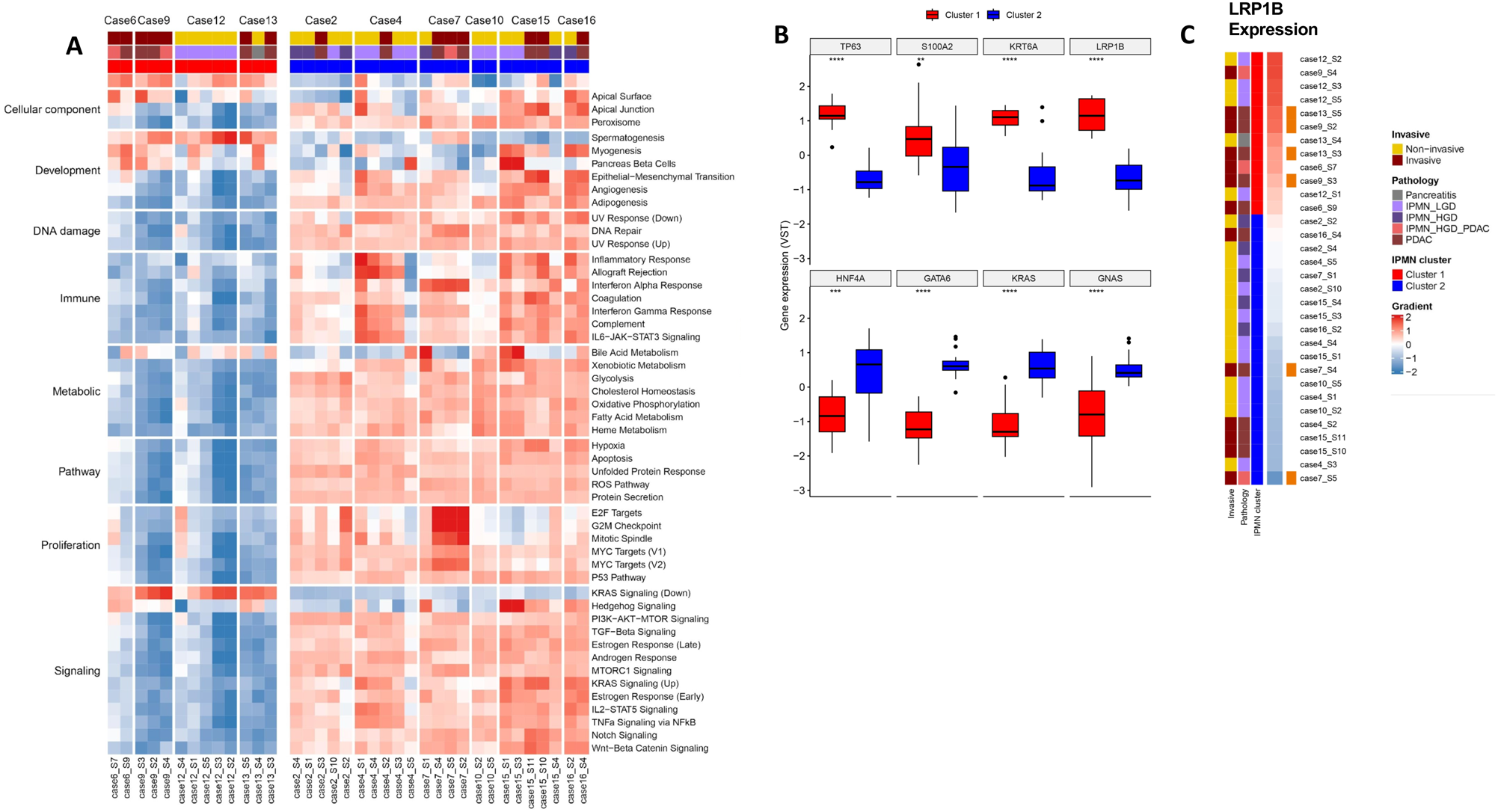
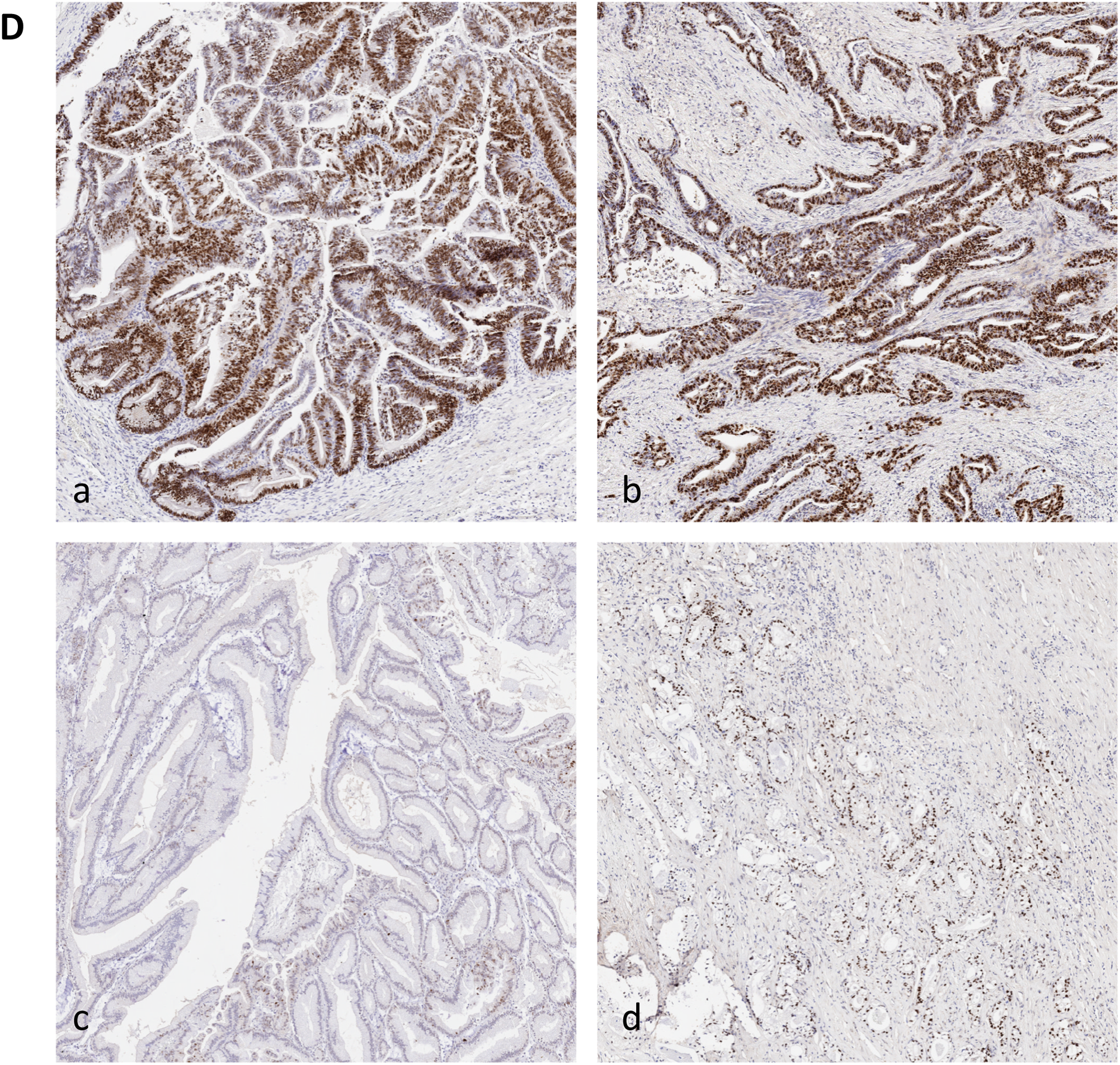
Gene and protein expression profiles across transcriptomic clusters. **A) Heatmap illustrating the differential enrichment of cancer-related pathways per sample within each case.** All samples from Cluster 1 are positioned on the left and those from Cluster 2 on the right highlighting distinct differences between the clusters as suggested by the PCA analysis in Figure 1. **B) Comparative Gene Expression Analysis between Cluster 1 and Cluster 2**. Differential expression of selected genes between the two transcriptomic clusters. Genes TP53, S100A2, KRT6A, and LRP1B exhibit higher expression levels in Cluster 1. In contrast, HNF4A, GATA6, KRAS, and GNAS show increased expression in Cluster 2. **C) Association of LRP1B Expression Levels with SNVs.** Relationship between LRP1B expression levels and SNVs (orange bars) in *LRP1B* across the samples. Samples from Cluster 1 are grouped at the top, demonstrating higher LRP1B expression, while samples from Cluster 2 are grouped at the bottom, indicating lower expression levels. **D) P53 immunohistochemistry on diagnostic FFPE samples.** Case 7 showed P53 mutation in both IPMN (a) and PDAC (b), while case 15 exhibited normal P53 expression in the IPMN (c) and P53 overexpression in the PDAC (d), validating the genomic results.

Gene expression was normalised, and a transcriptomic gradient score was assigned to each sample in the current cohort, as well as the previously published ICGC PACA-AU(19) and TCGA-PAAD cohorts(20), with a higher score indicating higher expression of genes associated with the previously described squamous molecular subtype (Figure 2A). This showed an overall trend of lower scores in the current cohort compared to the ICGC and TCGA PDAC cohorts, indicating higher prevalence of the Classical tumour subtype (Figure 1E), suggestive of more progenitor-like, earlier molecular stage disease with better prognosis. Uniform gradient scores of the Classical subtype were seen in Cases 2, 10 and 12, all characterised by non-invasive IPMN, reflecting overall stability within each tumour. Conversely, significant inter-sample variation within the same case was observed in Case 4, where one LGD sample (S1) exhibited Squamous subtype features as well as a higher number of SVs and CNAs compared to the other samples (S15, 6, 7 and 9) (Figure 2A).

To further interrogate the dynamics in the tumour microenvironment during IPMN progression, we employed EPIC cell proportion estimates(21), allowing comparative analyses among samples within each individual case (Figure 4A). Cluster 1, despite having higher Squamous subtype scores, demonstrated higher populations of CD8+ T cells (0.08 vs 0.01, *p* = 0.01), when compared to Cluster 2, whereas Cluster 2 exhibited higher populations of macrophages (0.006 vs 0.009, *p* = 0.04) (Figure 4B). We also observed that PDAC samples harbour significantly higher cancer-associated fibroblast (CAF) populations (median 0.24 vs 0.04, *p* = 0.04), and lower populations of uncharacterised epithelial cells than in premalignant IPMN (median 0.66 vs 0.52, *p* = 0.01), (Figure 4C, D) indicating increased stromal components associated with invasive PDACs. PDAC samples in general exhibited lower counts of CD4+ and CD8+ T cells compared to the matching IPMNs, such as in Cases 7, 9, 15, and 16. However, these differences were not statistically significant overall. IPMN-LGD samples such as those in Cases 10, 12 and 15 revealed uniformity in cellular composition among samples, with increased number of CD4+ T cells and reduced CAFs, compared to high-grade lesions. Using ESTIMATE scores(22), we summarised the stromal and immune components of each sample, confirming most PDAC samples with positive Stromal Scores except for Case 7. This case displayed histologically prominent inflammatory infiltration in all samples. Overall, higher Stromal Scores correlated with an increase in CAF population and enrichment of squamous subtype (Figure S6, S7, S9).

**Fig. 4.**
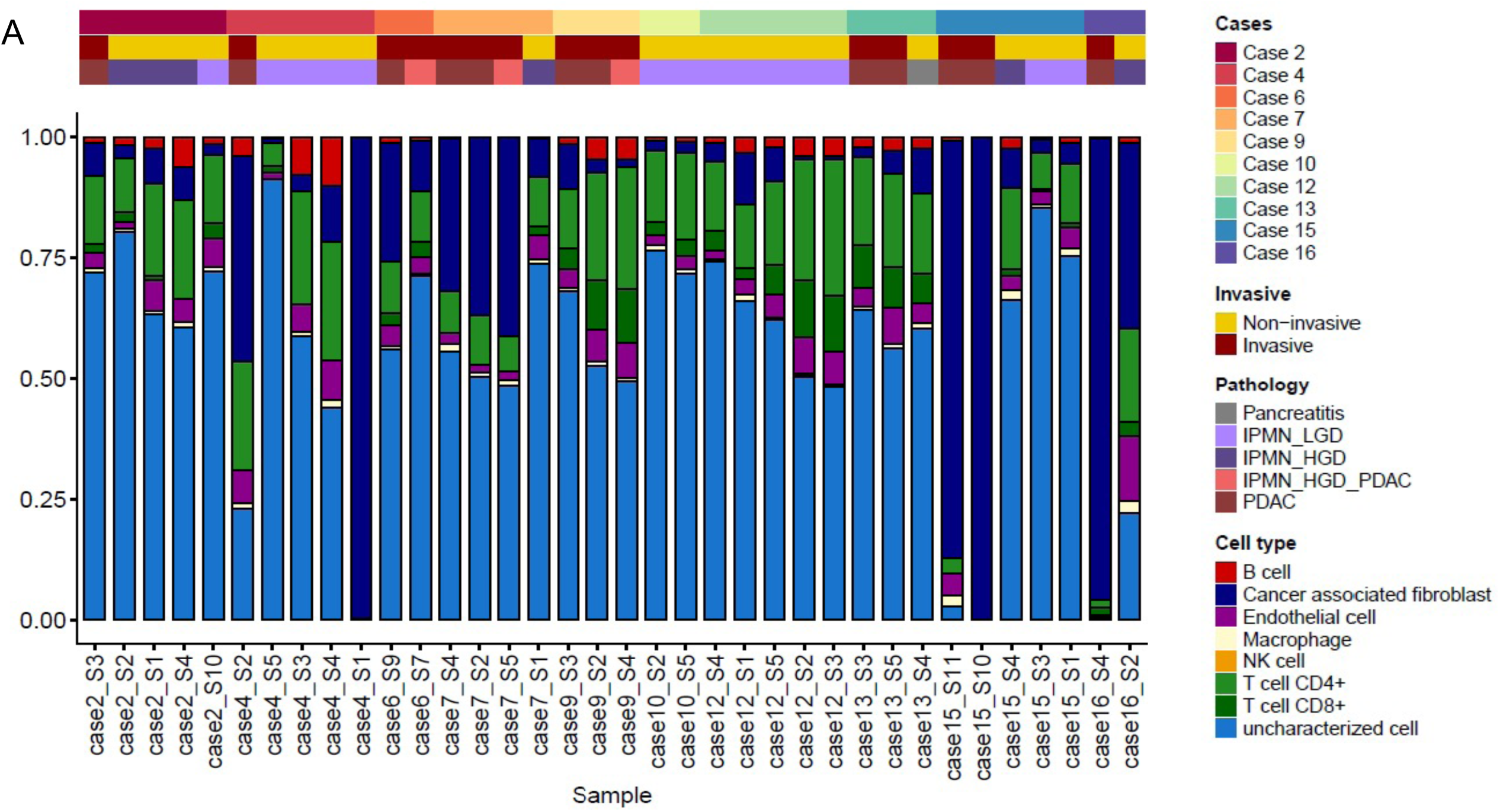
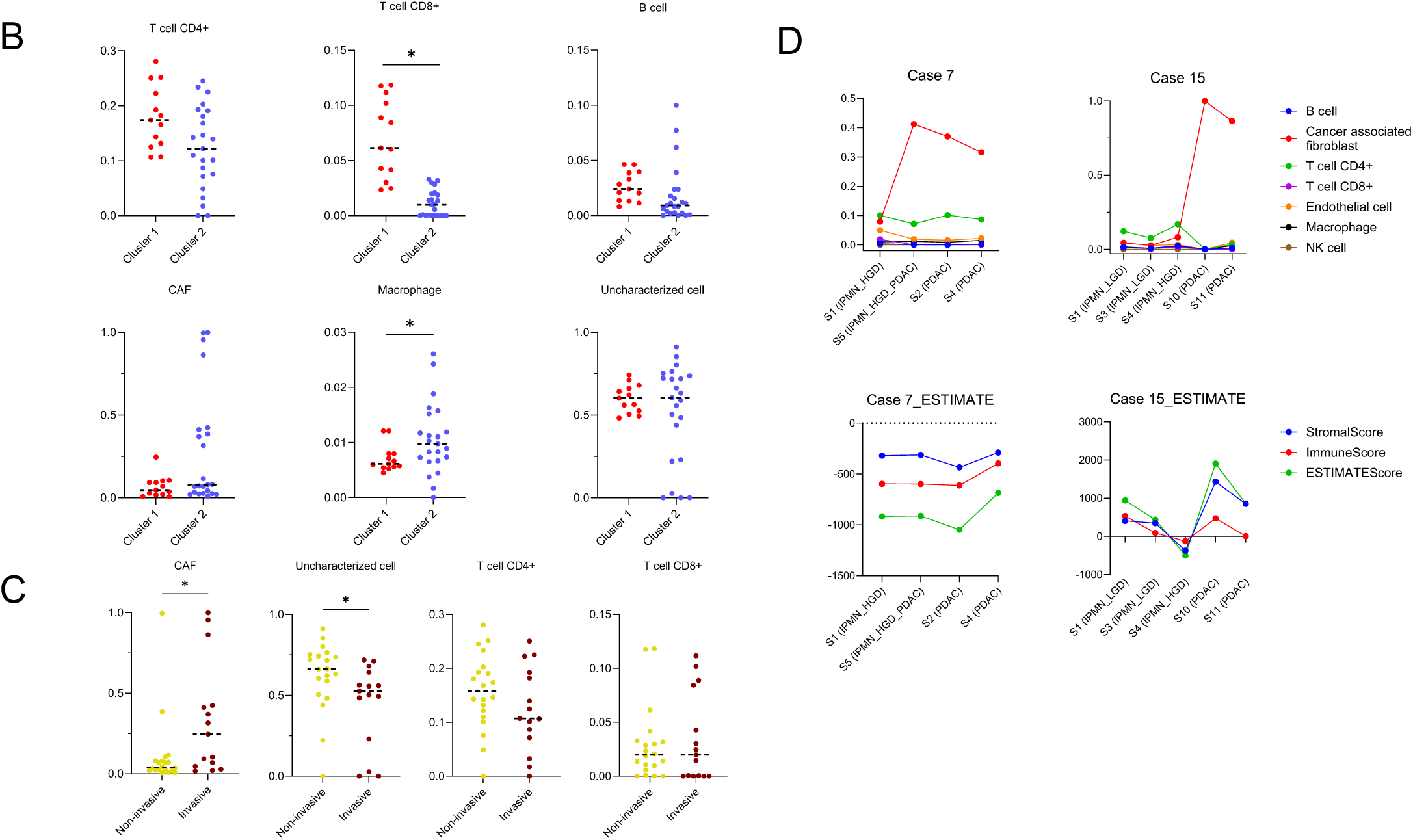
Deconvolution Analysis Revealed Different Cellular Composition Associated with IPMN Progression. A) EPIC cell proportion estimates for each sample, providing a comprehensive view of the cellular composition within each case. B) Comparison of cell proportion estimates between Cluster 1 and Cluster 2, revealing higher rates of T cell CD8+ in Cluster 1, and increase macrophages in Cluster 2. C) Representative cases (Case 7 and 15) illustrating the dynamic changes in immune cell composition and ESTIMATE-derived scores with disease progression from non-invasive IPMN to PDAC. D) Differences in cell proportion estimates between non-invasive and invasive samples, highlighting an increase in CAF and a reduction in uncharacterized cells, correlating with the transition from non-invasive to invasive stages.

## DISCUSSION

Here, we present the first analysis of WGS and RNAseq from multi-regional sampling of a cohort of patients undergoing surgical resection for IPMNs. These samples represent different grades of dysplasia and matching invasive cancer, enabling the identification of early molecular events contributing to malignant transformation of premalignant IPMN to invasive PDAC. Various classes of somatic mutations were observed in coding or non-coding regions in both IPMN and PDAC, with PDAC associated with a significantly higher number of simple and complex structural variations than IPMN. Despite sharing critical genomic alterations, such as *KRAS* mutations and *CDKN2A* losses, IPMNs and PDACs also exhibit distinct genomic profiles, for exmple SNV and Indels of *TP53* and *LRP1B*, losses of *RNF43* and *U2AF1* were all enriched in high-grade lesions, and *GNAS* mutations in low-grade lesions. *LRP1B* and *U2AF1* alterations emerge as potential novel drivers for IPMN-derived PDACs, in addition to the conventional PDAC related alterations previously described in both IPMN and PanIN precursors(23,24). These observations underscore a progression model characterised by an increasing accumulation of genomic alterations, including PDAC driver mutation and intricate chromosomal rearrangements(25).

We demonstrate two distinct evolution patterns, namely emergence of IPMNs and PDACs from a single ancestral origin, and parallel emergence of independent IPMN and PDAC tumours. The IPMN-only tumours are characterised by a relatively low mutational burden, suggesting their early appearance within each lesion. The acquisition of driver mutations plays a crucial role in the progression of one clonal trajectory over the others, leading to an accumulation of genomic aberrations and promoting the emergence of subclones. Notably, in cases 4 and 15, we identified distinct mutations in identical driver genes, in line with previous studies that deployed exome or targeted sequencing technology(7–9,24). This observation supports the hypothesis of convergent evolution, with mutations in the same gene across different subclones indicating parallel adaptation mechanisms. The absence of shared genomic alterations, in either structural or copy number, across these samples reinforces the theory of a non-genomic defect that initiates the tumorigenic process in the pancreatic ducts. Conversely, most cases presented a single genetic origin, representing a single ancestral cell giving rise to all subsequent tumours. A single MRCA initiated a lineage of one (Cases 3, 13 and 12) or more subclones (Cases 7, 9, 10 and 16), leading to the formation of HGD or PDAC lesions though the accumulation of additional genetic alterations over time. However, even in cases with a MRCA, we observed subclones present in IPMN-HGD samples that were absent in the corresponding PDAC, indicating parallel concurrent and independent genomic evolution of HGD alongside the PDAC, albeit from the same genetic origin. Previous WGS mutational signatures studies in PDAC defined 4 major classes, namely age related, double-strand-break-repair (DSBR), MMR, and signature 8(26,27). In the current study, the most common SNV mutational signatures, SBS1, 5, and 40, appeared early and tend to be maintained throughout the cancer cells’ lifetime, while Indel signatures, such as ID1 and ID2 are frequently present only at the early stages of evolution, suggesting that patients’ age and defects in DNA repair mechanisms may contribute to the initiation of tumourigenesis(13,14,28). Signatures of CNAs and SVs are involved in the critical processes during tumour evolution, as increased CNAs and SVs are seen in PDAC samples, with cases showing branching clonal evolution being characterised by a significantly higher level of genomic instability(4,15).

Transcriptomic analyses(6,29–31) demonstrated that PDAC is likely defined by two main broad molecular subtypes with various hybrids in between, characterised by differential prognosis and response to therapy(32). Transcriptomic analysis of the current cohort revealed distinct clusters of gene expression, which when integrated with the inferred phylogenetic trees, provided further understanding of the phenotypic characteristics among different samples. We observed changes in the microenvironmental composition during IPMN progression with an increase in CAF populations, and a decrease in epithelial cells in PDAC. *LRP1B* is a gene implicated in the WNT pathway like *RNF46*, and is frequently mutated in cancers such as lung, melanoma and gastrointestinal tumours(33–35), and is associated with higher TMB, increased immune infiltration, and improved responses to immunotherapy. In our study, Cluster 1 samples showed increased *LRP1B* mRNA expression, reduced Wnt pathway activity, and a higher number of SNVs. This Cluster is also characterised by greater TMB and enhanced immune infiltration, suggesting a link between genomic changes, RNA expression, and immune adaptation in the tumour microenvironment. However, the exact role of *LRP1B* as a driver in these processes remains to be fully elucidated. Our study is limited by a small number of patients despite multi-regional sampling, and likely does not capture the full spectrum of IPMN molecular variations. IPMNs are indeed characterised by vast heterogeneity in morphology and clinical behaviour and outcomes. These differences might underlie varied patterns of cancer evolution and progression, therefore requiring further validation in larger patient cohorts to enhance the robustness of our findings. Lastly, our reliance on bulk whole genome and RNAseq from frozen tissue samples does not allow for the spatial localisation of different genetic clones within FFPE tissue. Gaining such spatial context could offer additional understanding of the interaction dynamics between tumour clones and different immune cells, increasing our comprehension of tumour behaviour and progression pathways.

## METHODS

### Patient cohorts and multiregional samples collection

Primary tumour tissue samples were prospectively procured from patients undergoing initial resective surgery at Glasgow Royal Infirmary, Glasgow, UK, and at the Pancreas Institute of the University of Verona, Verona, Italy. The decision to proceed with surgical resection was based on the risk of the pancreatic cystic neoplasm harbouring a high-grade lesion (high grade of dysplasia and invasive cancer), in accordance with the most recent clinical guidelines for IPMN management(2,36,37).

Our multiregional sampling protocol was designed to collect samples that spanned all grades of dysplasia and invasive cancer within the same neoplastic lesion. Each pancreatic lesion was divided into 4 to 9 segments, with the relative positions of each segment meticulously recorded in a clockwise numeric order for spatial reconstruction, adhering to the institutional grossing protocol. During the grossing examination, care was taken to ensure that the collected samples were from segments of the same lesion with no normal tissue in between, to avoid the inclusion of concomitant but separate tumours. Blood, duodenum, or spleen samples were utilised as germline references.

All tissue samples were immediately snap-frozen in liquid nitrogen post-collection and stored at −80°C. DNA and RNA were extracted from bulk frozen tissues using an AllPrep DNA/RNA Kit (Qiagen), and from blood samples using a Nucleon Genomic Extraction kit (Gen-Probe), following the manufacturer’s instructions. DNA and RNA quantification and quality control were conducted using both Nanodrop and High Sensitivity Qubit (Thermo Fisher Scientific).

Hematoxylin and eosin-stained frozen sections with histologically confirmed IPMN or PDAC samples, as determined by an expert pathologist, and with an adequate amount of DNA and RNA, were deemed suitable for sequencing. All H/E slides were digitised for a second review by an expert pathologist and for cellular quantification.

Specific immunohistochemical staining on FFPE diagnostic slides of non-invasive IPMNs and the corresponding PDAC was obtained following standardized procedures(38–40), and evaluated as per manufacturers’ instructions. For all cases, the following antibodies were tested: P53 (clone: DO-7; 1:50 dilution; Novocastra/UK), SMAD4 (B-8; 1:1000; Santa Cruz/USA), and LRP1B (Polyclonal/rabbit; 1:100; Sigma-Aldrich/USA).

### Read alignment and somatic variant calling

Lanes of paired raw FASTQ reads from each sample were merged through fastqtobam tool and then aligned to human reference genome (hg38) using Sanger bwamem2 mapping workflow cgpWGS330 (ftp://ftp.sanger.ac.uk/pub/cancer/dockstore/human/GRCh38_hla_decoy_ebv/core_ref_GR Ch38_hla_decoy_ebv.tar.gz).

Somatic SNVs and InDels were identified from matched normal and tumour pairs using Sanger cgpwgs210 within a singularity (https://github.com/cancerit/dockstore-cgpwgs/wiki/Running-under-singularity) and Mutect2 from GATK 4.1.8 (https://gatk.broadinstitute.org/hc/en-us/sections/ 360009656231-4-1-8-0).

In order to generate VCF datasets for further analysis, mutational variants from these tools were selected with pass parameter and finally intersected using bcftools 1.11. In parallel, Breakpoint variants were isolated and assembled by cgpwgs210-BARSS. The SV files across all samples were filtered via SURVIVOR software and processed per sample using Python package Scikit-Allel to compute aggregated counts of each SV type: deletion, inversion, translocation, tandem duplication and multiple features. Fifty known drivers were queried from the final relevant datasets based on COSMIC cancer gene census(41,42).

### Detection of subclonal copy number alterations (SCNAs)

By inputting the alignments from normal-tumour pairs and Battenberg packaged reference (hg38), Battenberg algorithm based-pipeline was employed to identify copy number and estimate tumour purity and ploidy as previously described(12) (v2.2.9;https://github.com/Wedge-lab/battenberg). Among the CNAs found in the set of samples, regions altered at a significant high frequency were identified using GISTIC2(43) with the following modified parameters: -brlen 0.7 and -conf 0.99.

### Mutation and copy number signature profiling

Mutation signatures were analysed by using SigProfilerExtractor based on a non-negative matrix factorization (NNMF) framework (v0.0.5.77; https://github.com/AlexandrovLab/SigProfilerExtractor). Signatures from De novo extraction and decomposition were profiled as single-base substitution (SBS96), double-base substitution (DBS78) and small insertion-deletion (ID83)(14). Using ASCAT matrix generated from CNA datasets, copy number signatures were extracted by SigProfilerExtractor(15). Based on the updated known signatures as in COSMIC database, SigProflerAssignment was utilised to retrieve decomposed signatures.

In order to minimise NNMF artefact potentials, initial signature extraction was performed with all available samples simultaneously(44) and each single signature assignment per sample was finally determined by the probability matrix.

### Mutation clustering and phylogenetic tree reconstruction

To model subclonal structure and construct phylogenetic tree from multiple normal-tumour matched samples, we combined various genomic datasets and applied a number of bioinformatics tools. For each sample, mutation allele fractions (MAF) of SNV and Indel were prepared by using alleleCounter and vafCorrect. Together with copy number and cellularity, these outcomes were submitted for clustering mutations to their mutation copy number using a previously described Bayesian Dirichlet process (DPClustering)(16), (v2.2.8; https://github.com/Wedge-lab/dpclust/releases). This estimation per sample was extended into n dimensions for 12 IPMN-PDAC cohorts with n related samples, where the numbers of mutant reads obtained from multiple related samples were modelled as independent binomial distributions. Variants from each cohort with an upper CCF boundary above and below 1 were considered to be clonal and subclonal respectively(45).

To determine the most likely phylogenetic tree we applied the “pigeon-hole” principle (PHP) or ‘sum’ rule and ‘crossing rule’ to mutational clusters within individual samples(12,45,46). The sum rule asserts that if a subclone A is ancestral to both subclones B and C and if the summed CCFs of B and C exceed the CCF of A in any sample, the relationship between the subclones must be linear; whereas the crossing rule asserts that if the CCF of B is higher than the CCF of C in sample X and the CCF of B is lower than the CCF of C in sample Y, then B and C are constructed as separate branches of the phylogenetic tree(47).

### Annotation of the trees with mutations and signatures

To annotate each tree with oncogenic or putative oncogenic alterations, in-house programs were utilised to connect multiple genomic variants and identify specific significance, including SNV, Indels, SV, CNV, mutation signature and cluster assignment information from mutation caller, Batterberg, signature extractor and msDPClustering.

### RNAseq analysis

Sequencing quality of fastq files was assessed with FastQC (https://github.com/s-andrews/FastQC) (v 0.11.9) and files were processed with fastp(48) (v 0.21.0) using default settings. Quantification was performed against GRCh38 using Salmon(49) (v 1.4.0). Salmon quantification results were imported into a DESeqDataSet using DESeq2(50) (v 1.38.3). Transcripts were mapped to genes using EnsDb.Hsapiens.v86(51) (v 2.99.0). Read count data were filtered to retain only those with normalised counts >= 5 in at least 18 samples (18,409 genes retained). Reads were transformed using the DESeq2 ‘vst’ function. PCA was performed with plotPCA (DESeq2).

Scores were calculated for each sample for the following molecular subtypes: Bailey subtypes(19) (ADEX, Immunogenic, Pancreatic Progenitor, Squamous), Collisson subtypes(52) (Classical, Exocrine, Quasi-Mesenchymal), and Moffitt subtypes(53) (Basal, Classical). Two further subtype scores, ‘Squamous score’ and ‘Classical score’, were calculated for each sample using a signature derived from RNAseq data for squamous vs non-squamous samples from ICGC PACA-AU(19). A transcriptomic gradient score was therefore obtained by subtracting the classical score from the squamous score. Samples were also scored for ten previously defined gene programmes associated with Bailey subtypes. Scores were calculated for 50 Hallmark(18) gene sets from MSigDB. Scores were calculated using the singscore(54,55) package and the VST transformed counts.

Cell type proportions were estimated using the EPIC method as implemented in the immunedeconv(21) package using TPM counts. The ESTIMATE(22) method was used to calculate stromal, immune and overall ESTIMATE scores, implemented in the estimate (https://bioinformatics.mdanderson.org/estimate/rpackage.html) package.

### Data analysis

Pipeline running, data processing, statistical analysis and visualisation were mainly performed in R and Python programming languages based on High Performance Computing (HPC) cluster, The University of Manchester. To determine an association between lesion grade and the presence of mutational types we applied Fisher’s Exact Test for genomic alterations, and the Mann-Whitney Test for expression and deconvolution data across different tumor stages. Additionally, to compare the overall number of SNVs, Indels, SVs and TMB between IPMN and PDAC, we used Fisher-Pitman Permutation Test. To assess differences in cell count data between Clusters 1 and 2, median values from samples of each case were considered. A *p* value of less than 0.05 from the estimation was considered significant.

## Notes

### Competing Interest Statement

The authors have declared no competing interest.

### Summary of Updates

Increased quality of the main figures.

