## Supplementary Figures for "Clonal evolutionary analysis reveals patterns of malignant transformation in pancreatic cancer from Intraductal Papillary Mucinous IPMN Neoplasms (IPMN)"

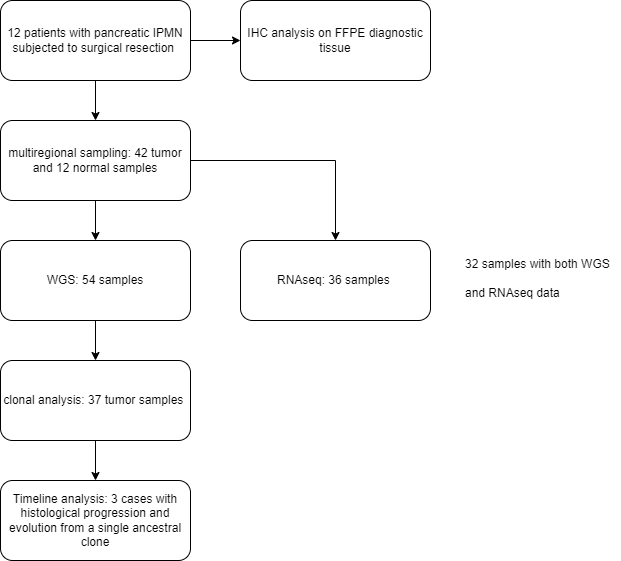
**Supplementary Figures**

**Figure S1A.** CONSORT diagram illustrating sample collection and analysis

**Figure S1B1 Comparison of Somatic Alterations Between IPMNs and IPMN-Derived PDACs.** Total 66,724 SNVs, 4683 Indels and 2447 chromosomal rearrangements in all samples, covering 48 drivers and 11200 genes

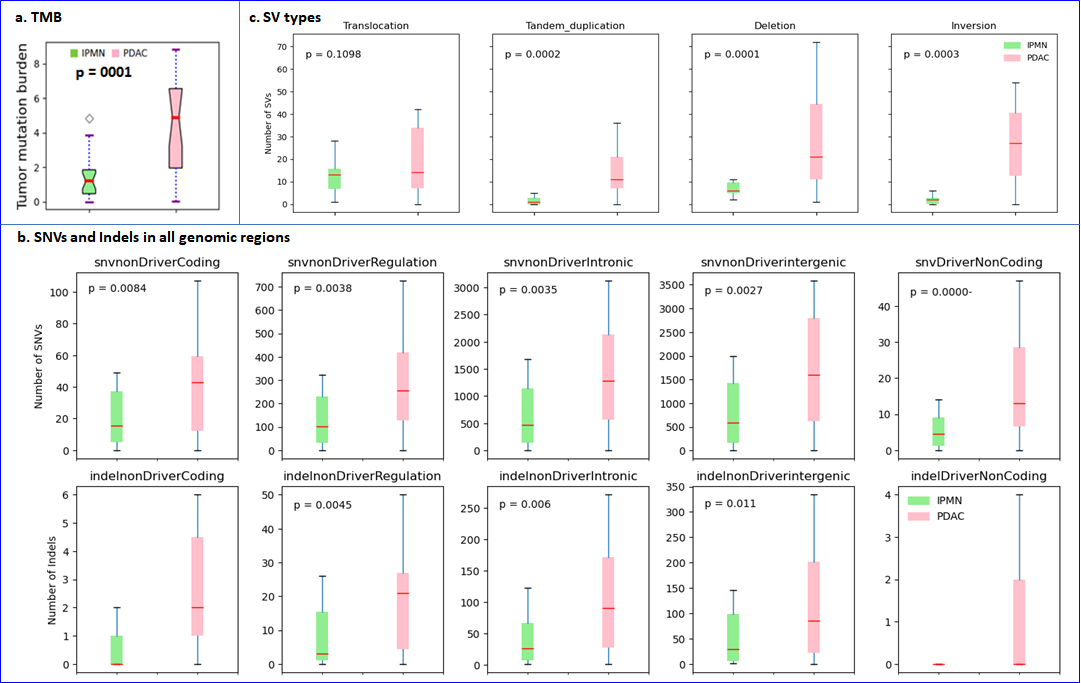

**Figure S1B2** Proportion of Mutational Signature in Each IPMN and IPMN-Derived PDAC

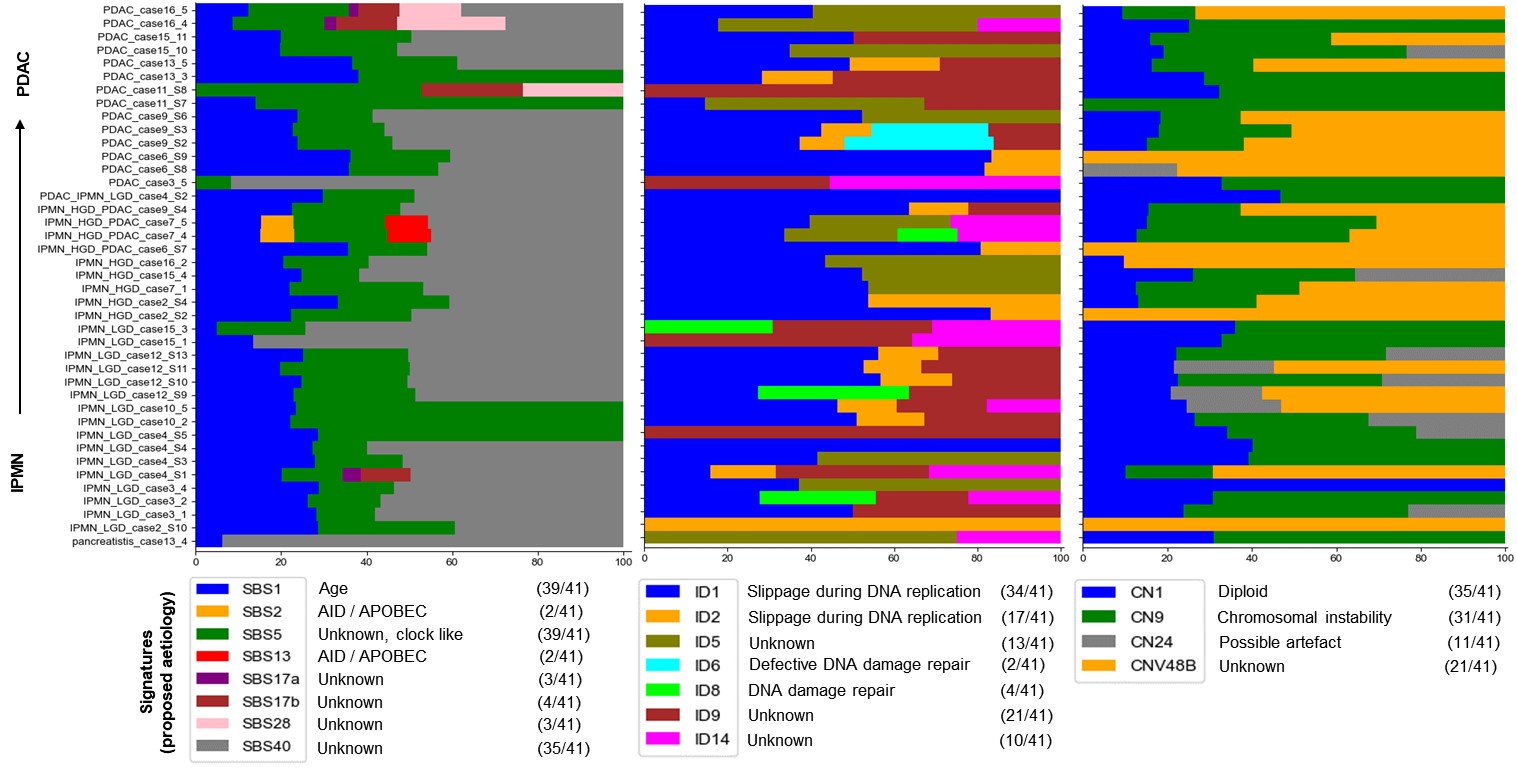

**Figure S1B3. Profile of the novel CNV signature CN48B**. This new CN48B is characterised by a large number of LOH segments with different lengths, which are distributed across dominant high copy numbers.

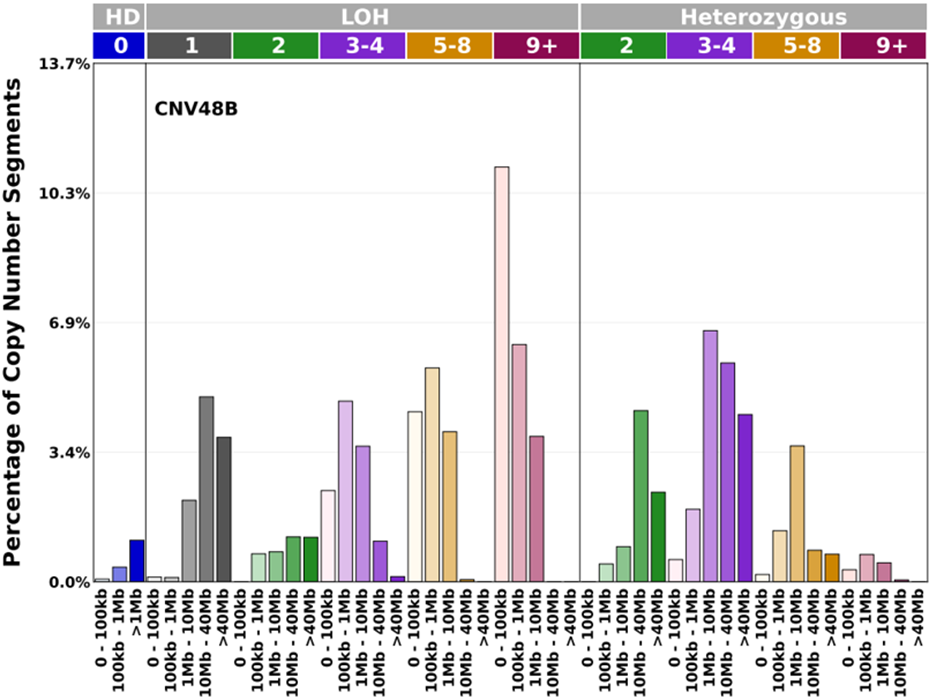

**Figure S1C1. Driver SNV / INDELs and CNV in IPMNs and IPMN-Derived PDACs (p < 0.05)**

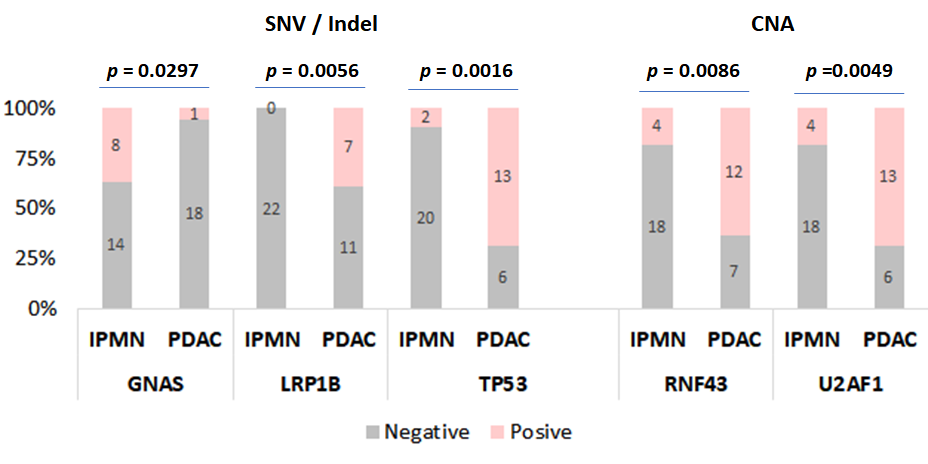

**S1C2. KRAS amino acid substitution (missense)**

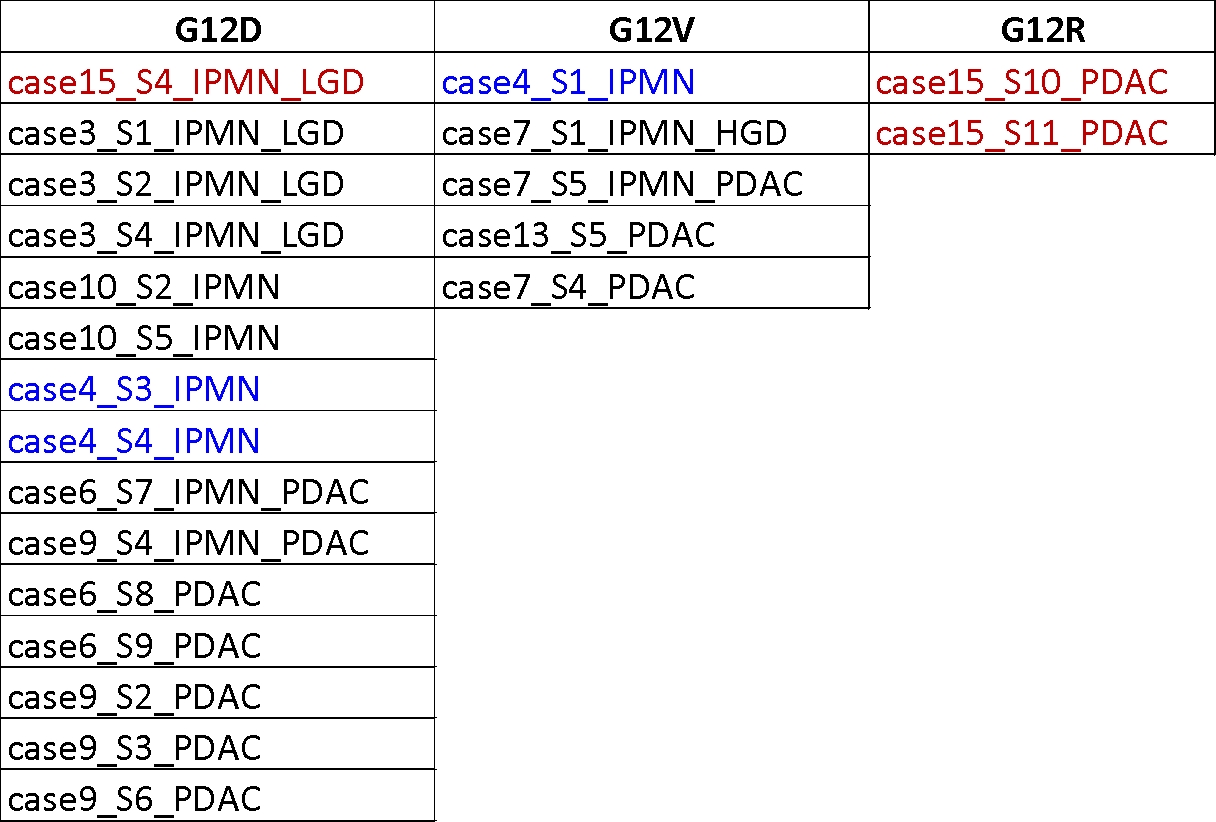

**Figure S1C3 Analysis of CCF and VAF in KRAS missense IPMN and IPMN-Derived PDACs**

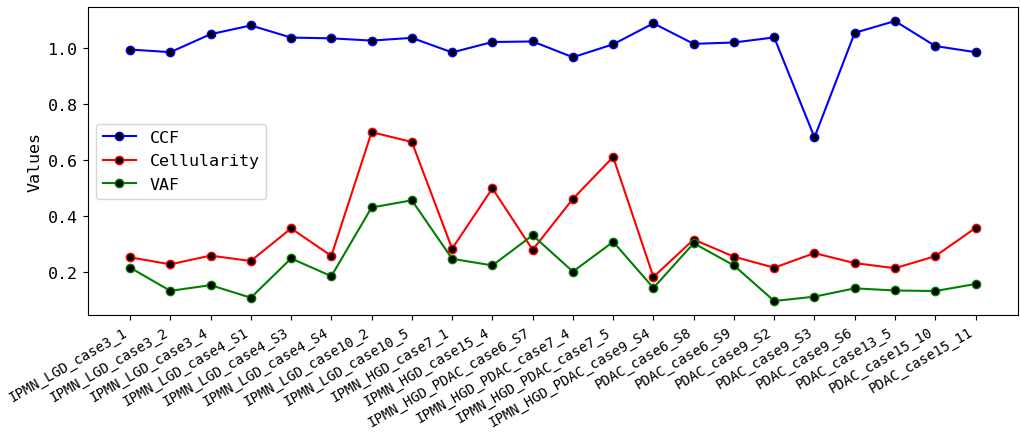

**Figure S2 Specific copy number alterations (CNAs) in** **IPMN and IPMN-Derived PDACs**

**B**

**A**

**B**

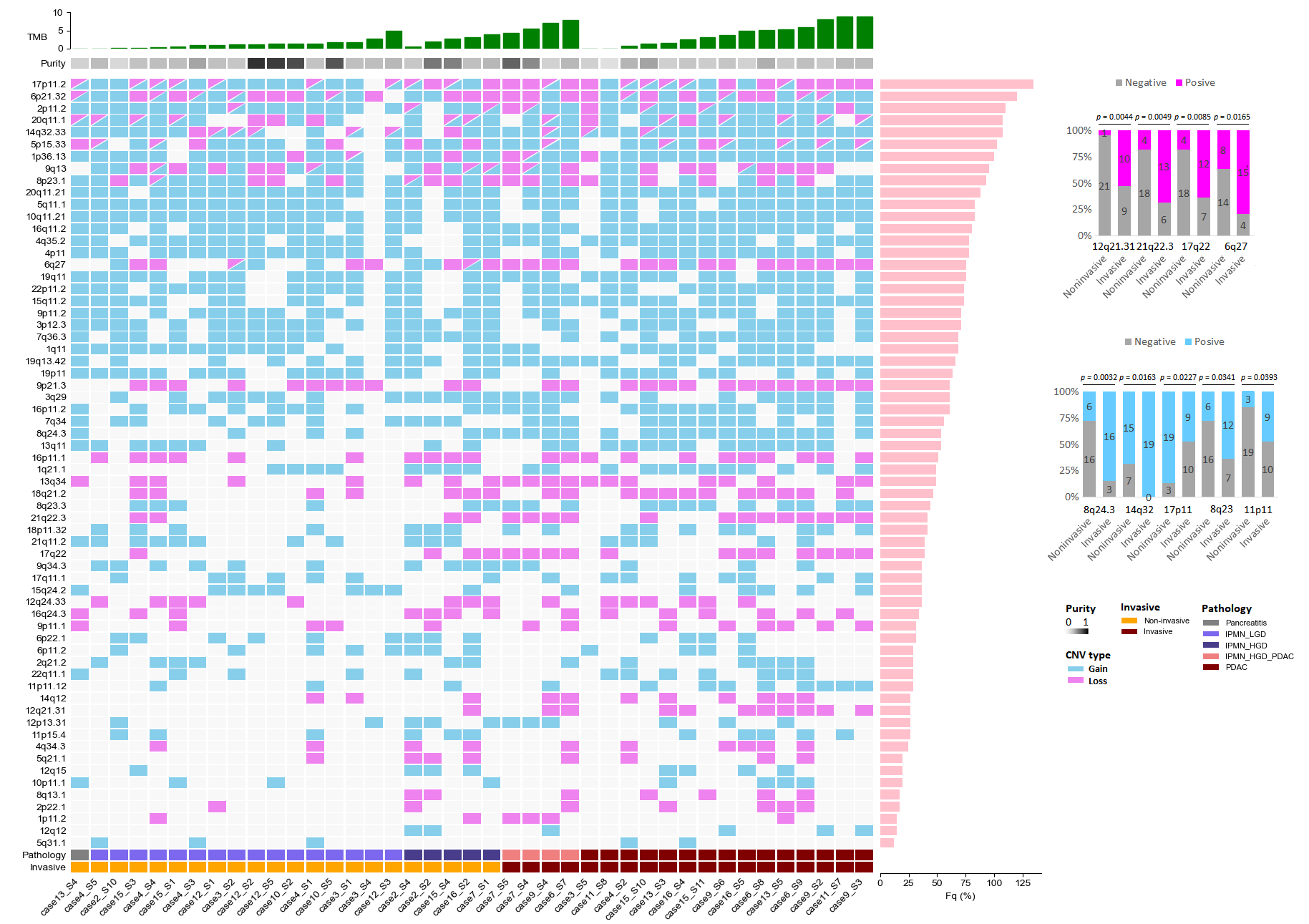

**Figure S3: Transcriptome sample correlation heatmaps.** The correlation heatmaps show the overall similarity of the expression profiles of samples. We calculated the Euclidean distance between all samples and presented by case.

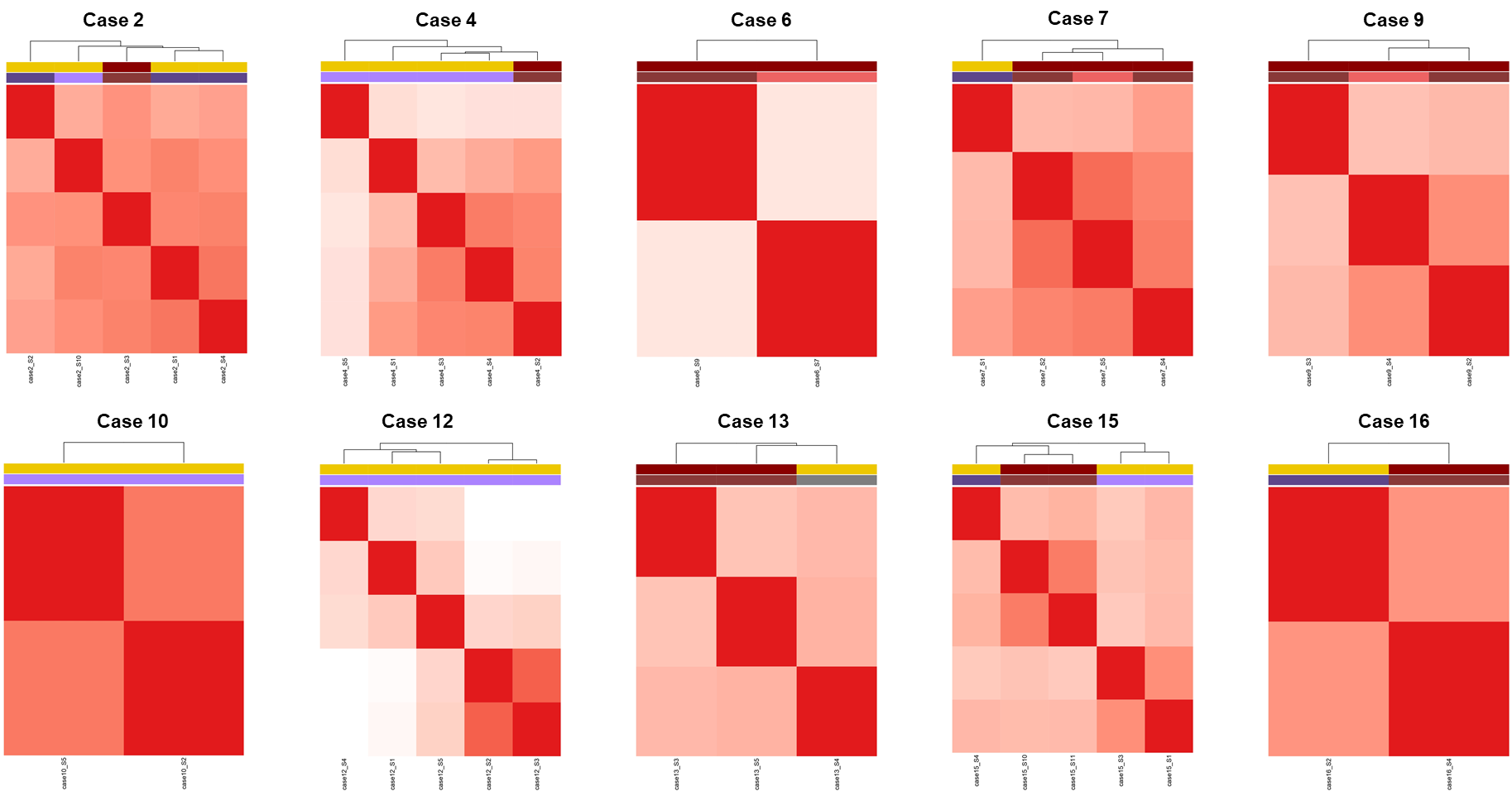

**Figure S4: Hallmarks of Cancer: Transcriptional networks dynamics among non-invasive and invasive samples (A) and among samples characterized by different histology within individual cases (B).** Metabolic pathways such as adipogenesis and fatty acid metabolism were generally more enriched in non-invasive samples, while glycolysis present higher expression in invasive samples within each case. The E2F targets and EMT pathways demonstrated clear differentiation in selected cases between non-invasive and invasive samples within single cases, with increased expression observed in high-grade lesions compared to the corresponding low-grade samples.

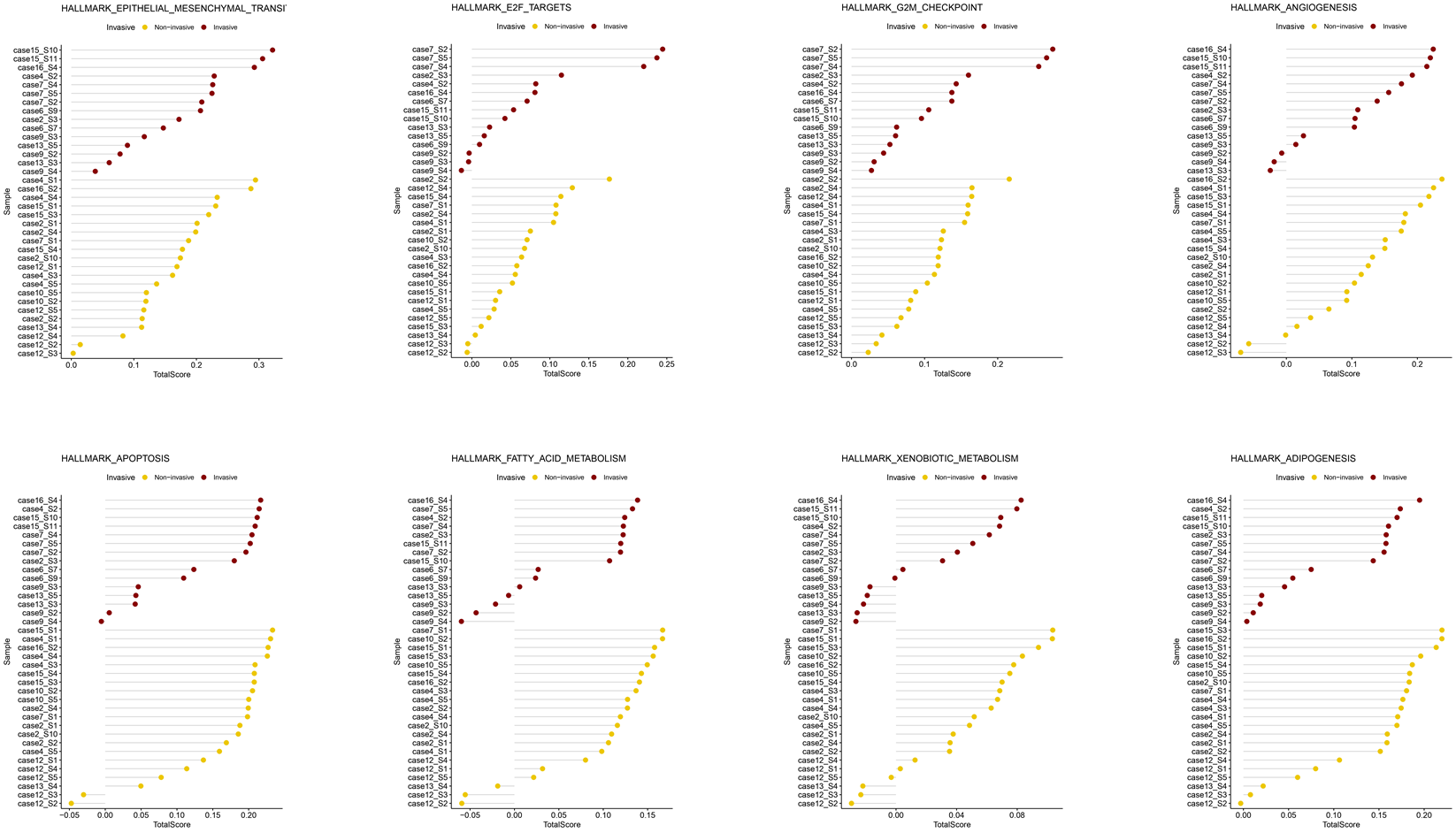
**A**

**B**

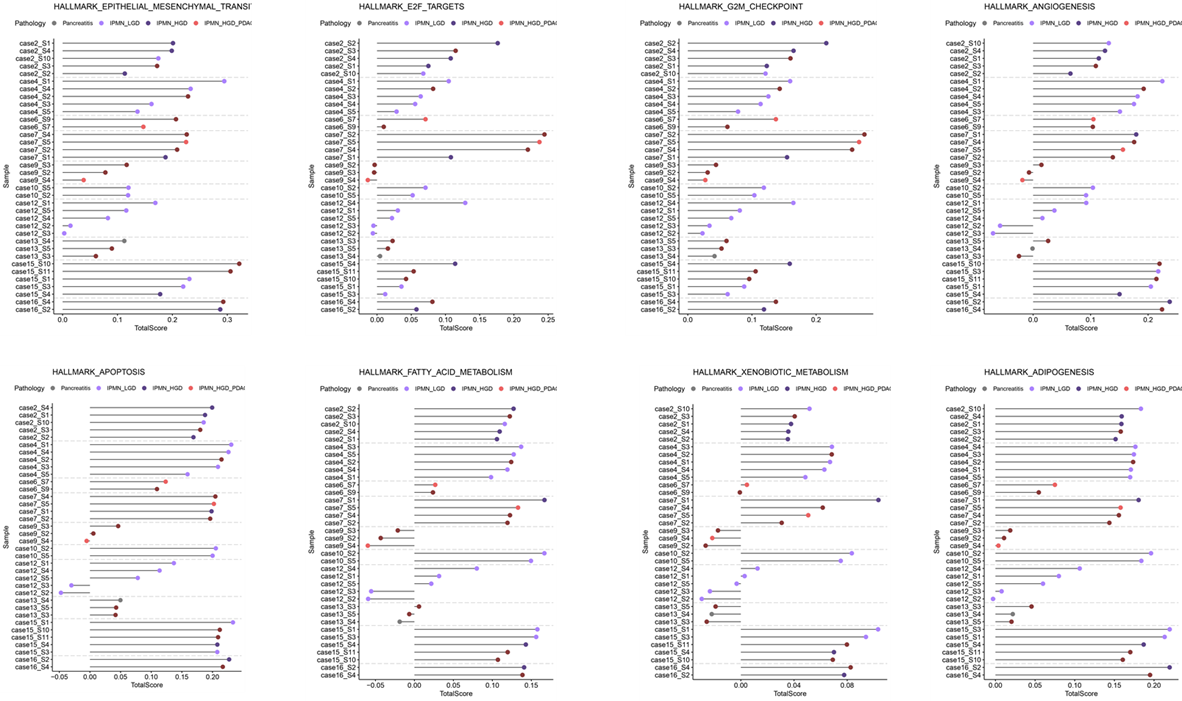

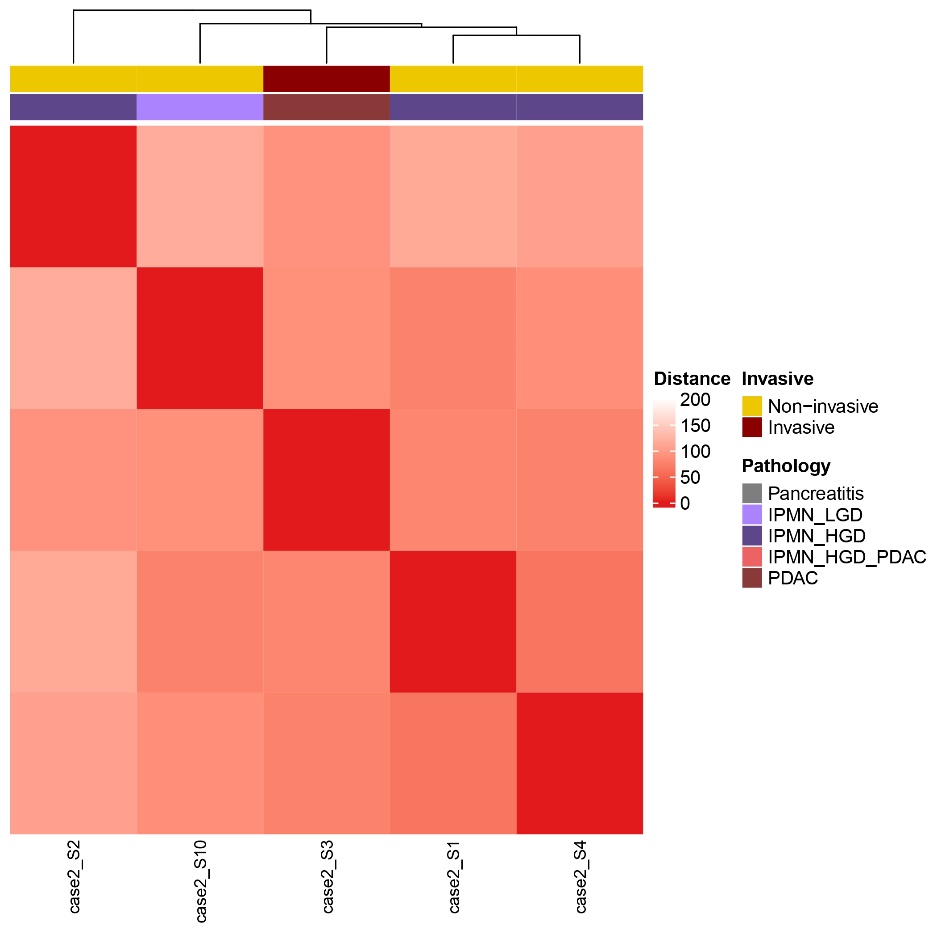

**Figure S5: PDAC transcription signatures**. Heatmap displaying PDAC transcription signatures for each sample, arranged from classical- (top) to squamous-related signatures (bottom) according to Bailey, Collisson and Moffit. The heatmap also includes gene programs significantly enriched according to Bailey et al. in the classical and squamous subtypes.

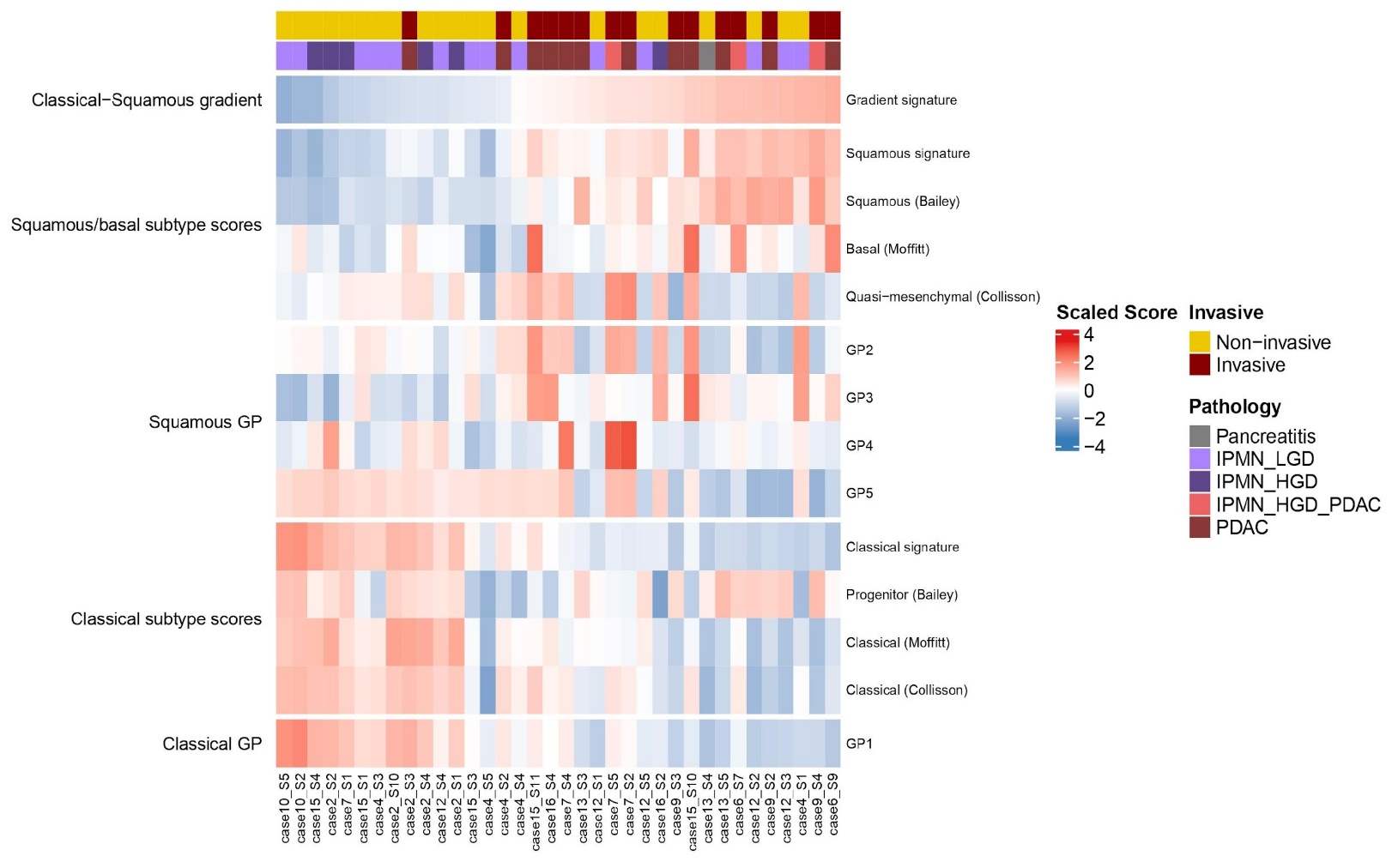

**Figure S6**

**Distribution of ESTIMATE, Stromal, and Immune Scores across samples.** Each sample is plotted based on its ESTIMATE score, arranged from the highest to the lowest. The corresponding Stromal and Immune scores for each sample are also depicted**.**

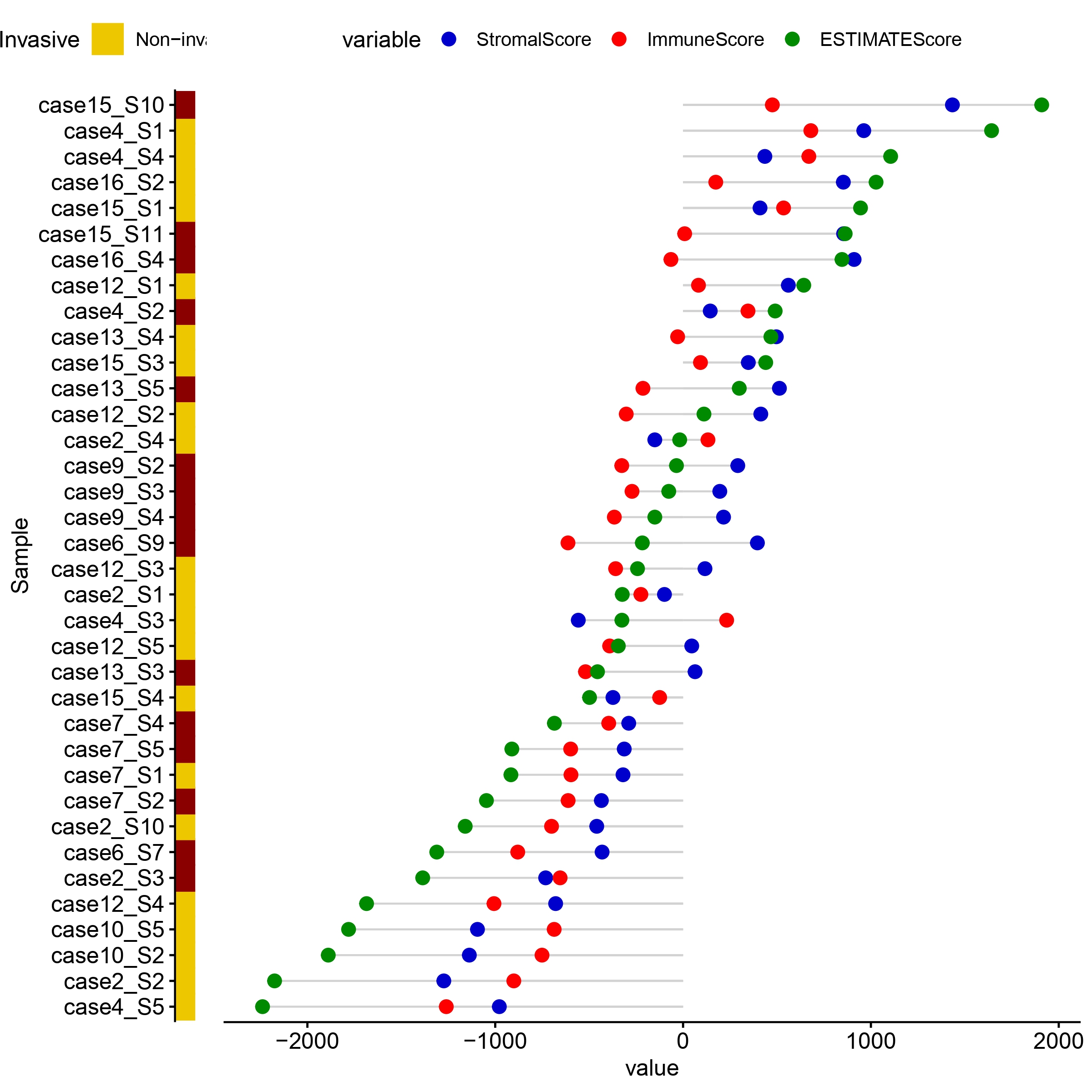

**Scatter plots illustrating the relationships between Squamous Score and the ESTIMATE-derived scores.** ESTIMATE (Estimation of STromal and Immune cells in MAlignant Tumor tissues using Expression data) is a computational tool designed to predict tumor purity and the presence of infiltrating stromal and immune cells in tumor tissues based on gene expression data. B shows a moderate positive correlation between the ESTIMATE Score and the Squamous Score, suggesting that as the overall tumor purity rises, the squamous characteristics might also intensify. C reveals a strong positive correlation between the Stromal Score and the Squamous Score, indicating that samples with a more pronounced stromal component tend to possess heightened squamous features. D displays a weak positive correlation between the Immune Score and the Squamous Score. While this relationship is not statistically significant, it suggests that the degree of immune infiltration in the tumor is not associated with the presence of squamous features.

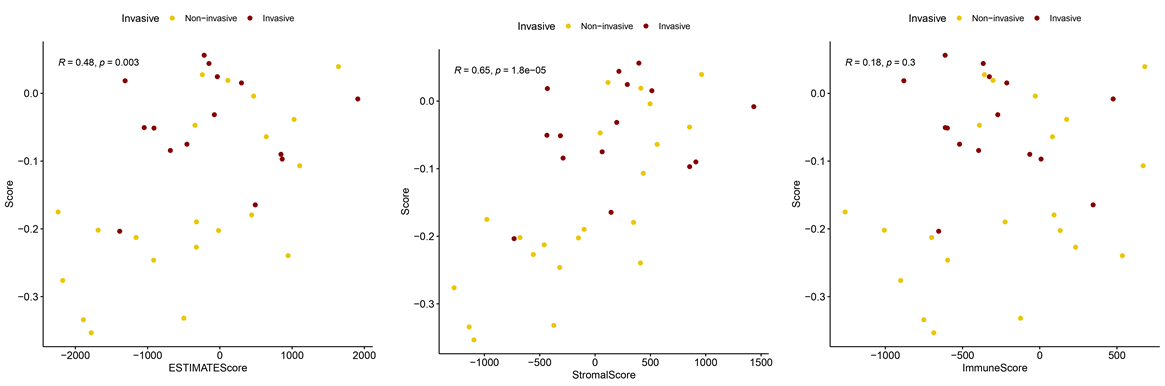

**Figure S7: Correlation Between ESTIMATE-derived Scores and Squamous Score in the ICGC Cohort.** The scatter plots display the relationships between the ESTIMATE-derived scores (x-axis) and the Squamous Score (y-axis) for 96 ICGC patients subjected to RNAseq. **A) Estimate score:** A positive correlation suggests that as the Estimate Score, which reflects overall tumor purity, increases, there's a tendency for the Squamous Score to also rise. **B) Stromal Score**: The stronger positive correlation here implies that samples with a more pronounced stromal component tend to possess heightened squamous features. **C) Immune Score:** The weak correlation suggests that the degree of immune infiltration in the tumor is not a strong determinant of squamous features. The distinct transcriptomic subtypes, as classified by Bailey et al., are highlighted, with the Squamous subtype distinctly segregated.

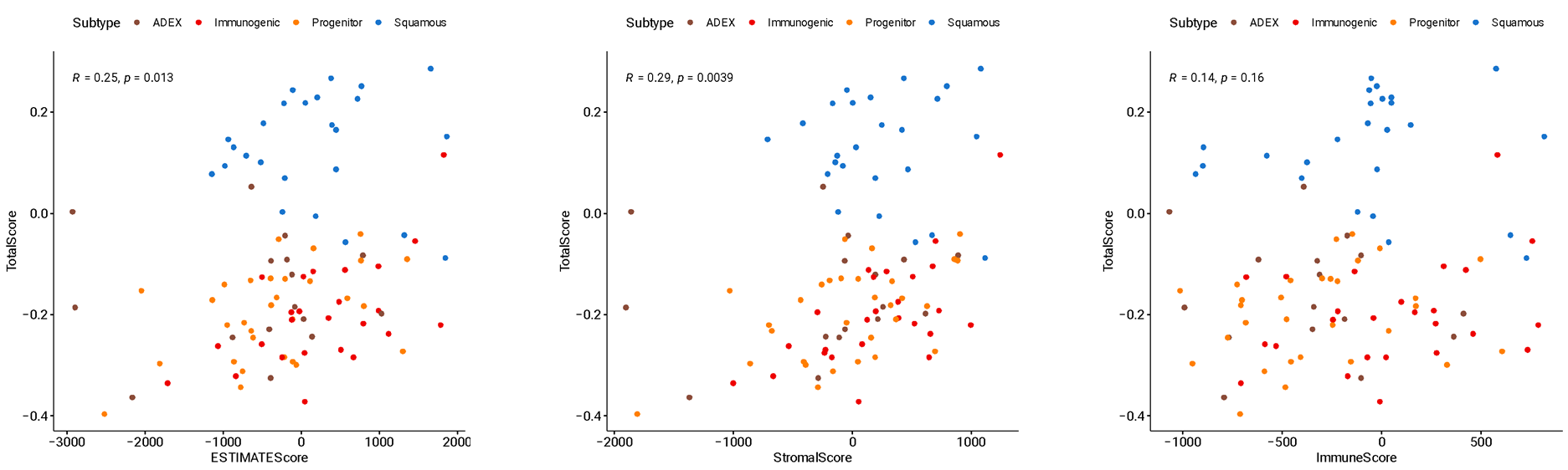

**Figure S8. Variations in Immune Cell Composition and ESTIMATE-derived Scores Across Cases.** This figure illustrates the dynamic differences in immune cell composition (excluding uncharacterized cells for clarity) and ESTIMATE-derived scores within each individual case.

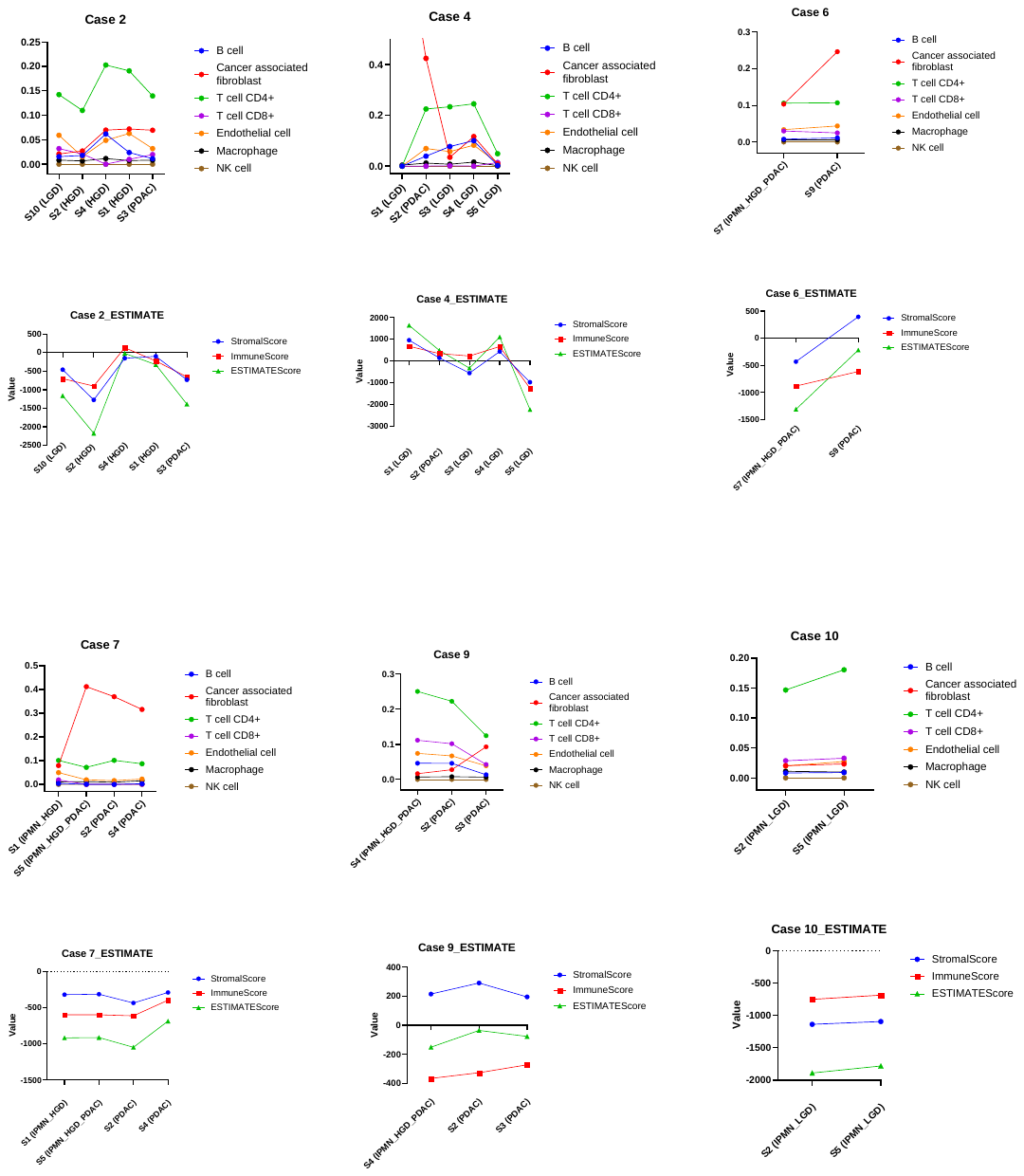

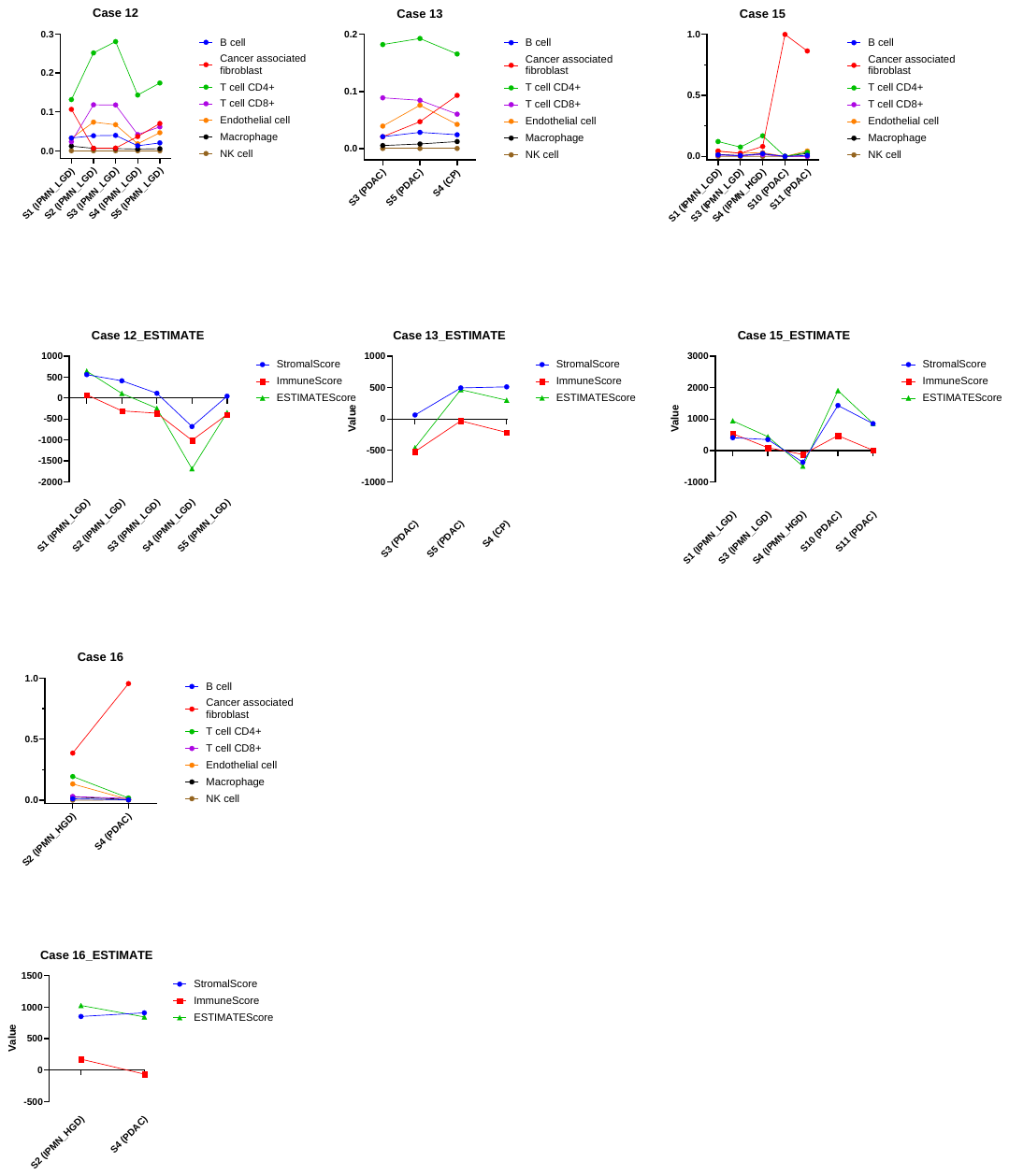
